# Contextual modulation of primary visual cortex by temporal predictability during motion extrapolation

**DOI:** 10.1101/2023.12.14.571621

**Authors:** Camila Silveira Agostino, Herman Hinrichs, Toemme Noesselt

## Abstract

Predicting future events is a fundamental cognitive ability which often depends on the volatility of the environment. Previous studies on apparent motion reported that when the brain is confronted with low levels of predictability, activity in low-level sensory areas is increased, including primary visual cortex. However, other studies on temporal predictability reported opposite effects potentially due to the influence of attention. It remains unclear, however, how temporal predictability modulates brain responses in a more ecologically valid real motion paradigm. Our study investigated whether motion extrapolation in high and low predictable contexts would differently modulate fMRI-responses in subject-specific primary visual cortex during visible and partially occluded stimulation. To this end, participants performed a modified version of the interception paradigm in visible and occluded phases, in which they observed a stimulus moving horizontally, then vertically at two different velocities. They were instructed to press when and where the stimulus would reach a given point-of-contact. In high predictable context, the velocity was identical during horizontal and vertical (occluded) movement; whereas, in low predictable context, the velocity could change during the vertical trajectory. MVPA results revealed accuracies above chance for all classification analyses carried out with low and highly predictable context data. Moreover, trial-history analysis showed that a change in trial type (constant velocity after change in velocity and vice versa) increased BOLD-responses in V1. This pattern of results suggests motion extrapolation can enhance activity in primary visual cortex, regardless of trial-specific predictability, but it is affected by recent trial history.

**Highlights:**

* Ignoring trial history high and low temporal predictability similarly enhance response in V1
* In recent trial-history analysis, low temporal predictability enhanced responses in V1 compared to high temporal predictability
* Shared regions inside primary visual cortex encode visible and partially occluded information

## 1. INTRODUCTION

Completing information partially missing from the environment – e.g. due to occlusion - is among the daily challenges that the cognitive system has to deal with, and for that, reliable predictive mechanisms are crucial. Prediction, as used here, can be understood as the ability of the brain to estimate future input based on (recent) past information (Alink et al. 2010). This idea that the brain uses internal estimates to make predictions about incoming events dates back to Helmholtz. Helmholtz proposed that to build a systematic representation of experiences, the mind uses mental adaptations referred to as “unconscious inferences” (Helmholtz 1866/1962; Patton, 2018), meaning that inferences, or predictions, affect directly our behaviour. More recently, Kahneman and Tversky (1972) extensively investigated how predictions reflect and affect behaviour and postulated that when making predictions and judgements under uncertain circumstances, humans seem to rely on heuristics – mental shortcuts – which may lead either to accurate judgements or systematic errors (Kahneman & Tversky, 1972; Tversky & Kahneman, 1971, 1973). In other words, prediction is an important mechanism used by the system not only to predict the future per se but to take into account the volatility, or the degree of unpredictability and variability, of the sensory environment.

In a highly volatile context, the sensory information may lead the system to less reliable predictions, hence larger prediction errors and smaller accuracy in sensory processing. A volatile environment here can be understood as unstable, fast-changing (Behrens et al., 2007). In learning paradigms, for instance, higher volatility resulted in higher learning rates: Behrens and colleagues (2007) investigated whether participants could keep track of statistical information presented in a reward environment and adapt their learning rate according to the observed changes. The authors demonstrated that participants were able to use prior and posterior accumulated information, even in highly volatile environment. In fact, volatility was an important factor used in the value judgement of each new incoming stimulus. In accord, Zylberberg and colleagues (2016) hypothesized that greater volatility would result in a quick diffusion of decision variable, i.e. decision reached after enough evidence has been accumulated. To test this hypothesis, humans and non-human primates were tested using a random-dot motion paradigm. The authors observed that, for both species, higher noise levels resulted in faster reaction time and more confident and accurate choices, suggesting that lower level of prediction can also aid information processing and decision.

Earlier studies developed theoretical models to explain how our brain works in a predictive fashion (Mumford, 1992; Rao and Ballard, 1999; Friston, 2003). One model which gained attention was the Predictive Coding model from Rao and Ballard (1999). In brief, the model assumes that different levels of a hierarchical model network make predictions and send them to lower levels via feedback connections, while higher levels receive back the information about the error between the prediction and the actual response via feedforward connections. This error is also used by the system to make corrections on the estimation of the input signal. This process would explain how predictions are coded in visual regions: lower visual areas learn statistical regularities from the environment and send forward the unpredicted aspics of the received input. Besides Rao and Ballard, others also attempted to mathematically explain how different systems works, by varying how the model is applied to the data and how the error is minimized (for review, see Spratling, 2017; Aitchison & Lengyel, 2017). Predictive Coding models have been used to test the mechanisms underlying temporal and spatial predictions in different domains, such as auditory (Baess et al., 2009; Heilbron & Chait, 2018, for review), motor (Shipp, Adams, & Friston, 2013), multisensory integration (Krala et al. 2019) and the visual domain.

In the visual domain, Alink and colleagues (2010) directly tested the predictive coding model by investigating whether highly predictable moving visual stimulus decreases activity in the primary visual area and whether the stimulus predictability also influences regions which send feedback information to V1, such as hMT/V5+ (Vetter et al., 2015). The authors used an apparent motion task and a random dot motion paradigm to test how unpredictable temporal and spatial information, respectively, increased response signal in V1. For apparent motion, highly predictable stimulus onsets indeed reduced activity in V1, compared to unexpected delayed onsets. For random dot motion, they also observed a decrease in response in V1, as well as in hMT/V5+, when the random dot motion direction was predictable. These findings supported the predictive coding model by showing that temporal and spatial predictable visual stimulations reduce activity in V1, while unpredictable visual stimulation enhances activity in this region. Later, Schellekens and colleagues (2016) extended these findings to V2 and V3, by testing how predictable and unpredictable contrast changes can modulate neural response in low-level visual regions, using random-dot motion at specific locations. Results indicated that when new dots entered the visual field by being presented on new locations, higher responses were registered in V1, V2 and V3 compared to dots which were already displayed in these areas. A recent EEG-study, which investigated the representation of real-time position of random moving dots, reported that such representations were updated in real time as the dots moved (Johnson et al, 2023), suggesting that V1 is able to update position information continuously. These findings are also in line with similar studies in humans (Ekman et al, 2017), which found anticipatory V1 responses for upcoming predicted events.

Using more complex stimuli Fischer and colleagues (2013) investigated how temporal predictability affects neural processing in low-level visual areas, during the categorization of predictable and unpredictable moving faces, when primed by an auditory alerting signal. One group of the participants was asked to judge whether the presented moving face was male or female, while the other group judged the direction of the stimuli. In half of the trials, the alerting cue was presented before the visual stimulus. Results indicated that participants’ behavioural performance was higher when the visual stimulus onset was predictable and when the alerting signal was coherent with it. However, a negative correlation was found between activity in V1 and the alerting signal, i.e. the larger the effect of the alerting signal in behavioural performance, the stronger the reduction in the primary visual cortex. These findings suggest, that the increase of temporal predictability reduced the BOLD signal in V1, corroborating previous studies and supporting the importance of temporal information in the prediction of future event.

In contrast to the above-mentioned findings, Coull and colleagues (2008) reported results apparently contradicting the postulates of the predictive coding model. Using a time-to-contact (TTC) paradigm, the authors investigated how fMRI-responses in low-level visual areas were modulated by temporal predictability during egocentric (subjective viewpoint) and allocentric (external viewpoint) viewing. In an ecologically valid driving simulation, participants were instructed to predict whether a car would touch a wall in one task, and in another task whether the colour of the car and the wall matched. It was hypothesized that proper attentional allocation would enhance stimulus detection, considering that spatiotemporal predictability is implicitly related to object-motion TTC task, while temporal predictability is explicitly associated with temporal cueing task. The authors observed increased fMRI-responses in V1 for TTC prediction during the egocentric judgements, likewise a variation in activity as a function of increasing certainty of collision; while allocentric judgements selectively enhanced responses in V5.

These findings were interpreted as an effect of attentionally-induced modulation in activity in V1. Moreover, this interpretation is in line with the findings of another electrophysiological study, which investigated how temporal and spatial information during dynamically occluded stimulation can expand attentional resources applied during the occlusion period (Doherty et al., 2005). The authors observed that the P1 component, which represent accumulated activity of many ventral and dorsal extrastriate visual regions (Di Russo et al., 2002; Foxe et al., 2005), was enhanced when spatial and temporal expectations were combined, suggesting that temporal predictability of occluded target plays an important role in establishing the efficacy of sensory processing. Moreover, they observed that spatial and temporal predictability work synergistically together and this interaction may affect the attention allocation to the reappearance of the occluded moving stimulus (Doherty et al., 2005).

Some researchers have tried to reconcile the opposing views generated by those studies which found evidence that predictability reduces response in the primary visual cortex, with those which found the opposite pattern. For instance, Kok and colleagues (2012) tested two hypotheses related to this contraction: (1) attention and prediction have an opposing relationship, meaning that the excitatory attentional effect outweighs the inhibitory effect related to prediction by enhancing activity when predicted attended stimuli are presented compared to unattended predicted stimulation; (2) attention and prediction have an interactive relationship, meaning that attention boost the precision of prediction, resulting in an enhanced activity promoted by the predictive error. The authors tested these hypotheses by measuring fMRI-responses in bilateral low-level visual areas (attended and unattended sides), while presenting cues which indicated the likelihood of the side where a Gabor patch would appear. Participants were instructed to indicate the orientation of the Gabor patch on the attended side and to ignore Gabor patches on the unattended side. Results indicated that reduced responses in V1 were observed on the unattended side when they were expected there; compared to enhanced responses in V1, V2 and V3 on the attended side for expected stimuli. The study provided evidence supporting the first hypothesis, which suggested that attention cancels the response reduction during high predictability, as the excitatory response related to attention enhances activity in low-level visual areas, but only on the attended side. Recently, we showed that when participants are asked to estimate the end positions of a partially occluded traveling stimulus, responses in V1 were enhanced along the occluded trajectory. Remarkably, even though the end position was always associated with a specific velocity, hence predictable, the participant’s engagement during the occlusion phase increased activity in V1 (Agostino et al, 2023). Note, that the trajectory was not straight but complex with a 90° turning point, which may have added an extra challenge to the visual processing and an extra strain on the attentional resources.

In this study, we aim at extending these findings by probing the influence on temporal stimulus predictability in the context of real motion extrapolation. To this end, we investigated whether dynamically occluded stimulation presented in high predictable (HP) and low predictable (LP) contexts would differently modulate fMRI-signals in V1, by manipulating temporal information. We hypothesized that, according to the predictive coding theory, signal in this region should be smaller during stimulus presentation in the HP context, which requires less attention compared to the more volatile LP context. To test these hypotheses, we used an adapted version of the interruption paradigm (IP - Battaglini & Ghiani, 2021), monitored brain activity using fMRI and acquired retinotopic maps, in order to identify subject-specific regions of interest. In addition to univariate analysis, multivariate pattern analyses were carried out in right upper and lower V1 (and right hMT/V5+, see supplementary material), as the stimulation was presented only in the left hemisphere, in order to investigate changes in representational pattern of activity in these regions (see Agostino et al., 2023 for a similar approach).

## 2. MATERIALS & METHODS

### 2.1. Participants

Eighteen participants (mean age 25.5, ±4.23, eight women) with normal or corrected-to-normal vision, no history of psychiatric or neurological disorders and no regular intake of medication known to interact with the central nervous system were recruited from the student community of Otto-von-Guericke Universität Magdeburg and gave informed consent to participate in the study according to the local ethics. In a two-day experiment, participants were exposed to eight tasks, four each day, while their brain activity was monitored by fMRI: two training phases (visible stimulation) and two tests phases (occluded stimulation). On a third day participants returned for a retinotopic mapping measurement. Volunteers were rewarded with 10 euros/hour or experiment credits. Two participants were excluded due to incomplete data acquisition.

### 2.2. Task

#### High Predictable Context - Visible Phase (training)

On a black screen, a white dot moved continuously horizontally (200 px) from the left side to the centre, then vertically upwards (200 px) or downwards (200 px) until it crossed a “X” mark (+120 px) (see Fig.1A). The vertical direction would be determined by the velocity of the horizontal movement, which could be 16.6°/s (fast, 0.250 s; 0.150 s after crossing the “X” mark) or 14.4°/sec (slow, 0.450 s; 0.266 s after crossing “X” mark). [Note that the velocities (or time displacement) used until the stimulus reached the “X” mark were the intervals used for modelling and comparison with participants’ response]. Hence, participants could easily learn the dependency of trajectory change on velocity since velocity was constant during horizontal and vertical trajectories. The participants were asked to keep their eyes on the fixation cross while attending to the sequence of movements and to indicate where and when the dot would end. We asked them explicitly to avoid pressing after the stimulus crossed the marks. Subjects responded by pressing the left button with their index finger or the right button with the middle finger in case the dot reached the upper “X” mark or lower one, respectively. Thirty trials of each condition were presented in four runs, i.e. 240 trials total (two runs with the configuration: up-fast/down-slow; and two runs: up-slow/down-fast). Intertrial interval was chosen from a Poisson distribution with values between 2 and 6 seconds and each run lasted around 8 minutes.

**Figure 1.**
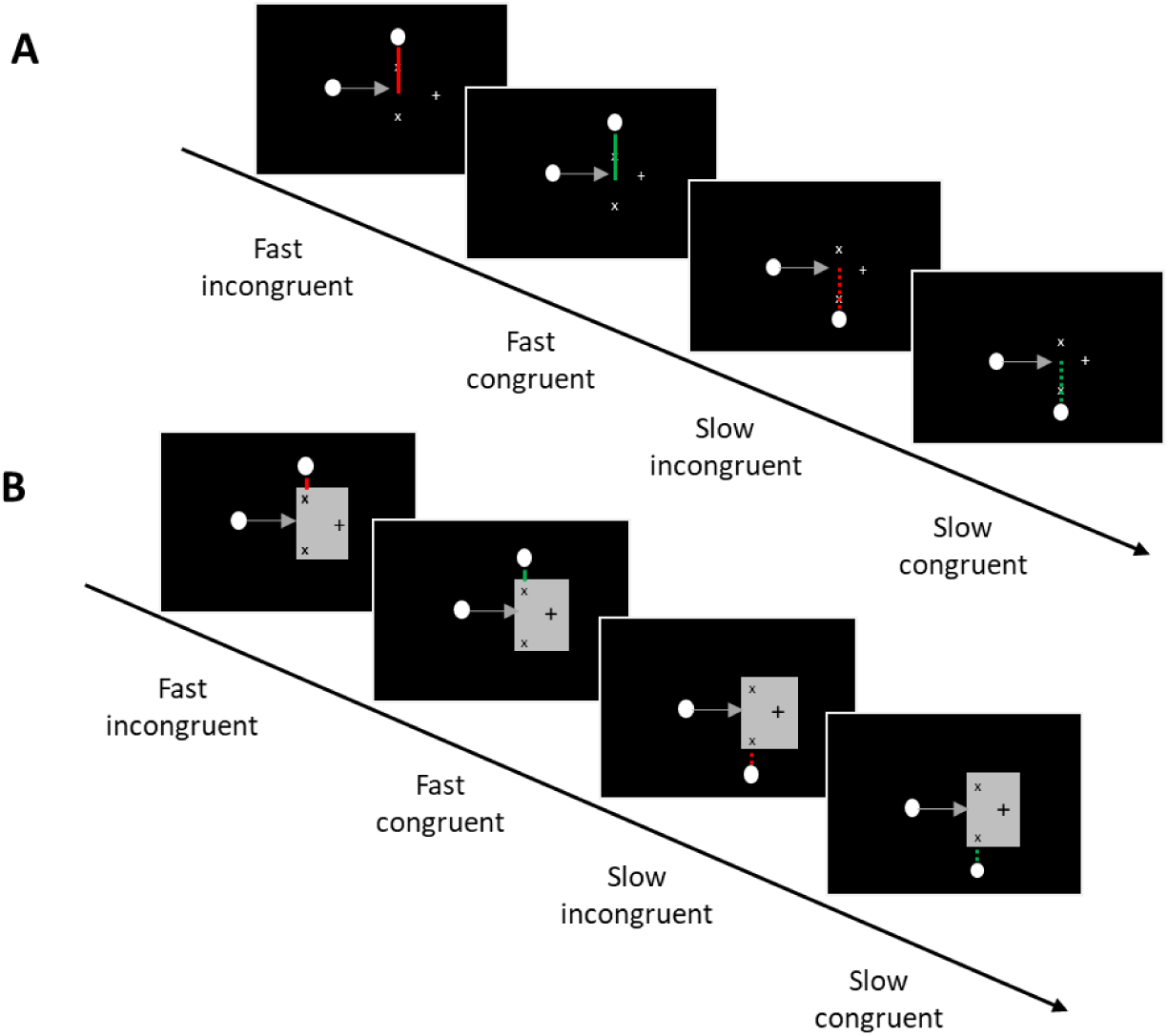
Experimental Paradigm: The full and dotted grey and red lines (not presented during the experiment) illustrate the two velocities (full = fast; dotted = slow). Different colours represent velocity changes (red = incongruent; green = congruent). The full red line represents fast incongruent trials, in which the stimulus travelled the horizontal trajectory with the fast velocity, but would slow down along the vertical trajectory. The dotted red line represents slow incongruent trials, in which the stimulus travelled the horizontal trajectory with the slow velocity, but would speeded up in the vertical trajectory (A) Visible Phase. Participants observed a white dot moving fast upwards or slow downwards (or vice-versa) and were instructed to estimate when the dot would reach the upper or lower “x” mark, based on the observed velocity. LP contexts contained the same stimuli as the HP context, but in 30% of the trials, the velocity-direction pairing would be incongruent, i.e. the dot changed speed at the upward/downward turning point (B) Occluded Phase. During the occlusion phase, participants performed the exact same task as during the visible phase, with the difference that the vertical trajectory was occluded by a grey rectangle. The dot reappeared from behind the occluder, so participants had feedback about the actual velocity. In the HP context, the velocity of the stimulus during the horizontal and vertical trajectory was always identical. Participants were explicitly instructed to respond when the target would reach the mark on the occlusion phase, i.e. before the reappearance of the stimulus.

#### High Predictable Context - Occluded Phase (test)

Our adaptation of the IP task was very similar to the visible task described above, with the difference that a grey rectangle, occluding the vertical trajectory, was presented during the whole trial. Participants extracted the temporal information from the horizontal trajectory, in order to make the judgement about the stimulus destination and time of contact (Fig. 1B). Once more, participants were explicitly instructed to avoid pressing the response button after the stimulus crossed the marks, i.e. reappeared. Fifty trials of each condition were presented in four runs, resulting in 400 trials (same configuration as above). Intertrial interval was chosen from Poisson distribution with values between 2 and 6 seconds and each run lasted around 13 minutes (Hinrichs et al., 2000).

#### Low Predictable Context - Visible Phase (training)

In this session, we manipulated the task predictability by introducing different probabilities of the temporal information. To this end, the paradigm in this phase remained the same as described above with the following exceptions, if the stimulus moved slowly along the horizontal trajectory, it could change speed in 30% of the trials and move faster along the vertical trajectory, or vice-versa. Seventy-two out of 240 trials were presented with the incongruent displacement time and the number of trials were counterbalanced across conditions. Note that the overall duration of speeding up incongruent trials and slowing down incongruent trials in the vertical trajectory was kept constant (700 ms), until the point-of-contact.

#### Low Predictable Context – Occluded Phase (test)

The task for this phase remained the same as described in the test phase of the high predictable context with the changes in velocity changes already reported above in the last section (visible phase of the low predictable context). The challenge in this phase was to estimate the displacement time without having the visual information of the vertical trajectory. The stimulus reappearance from behind the occluder served as feedback, but participants were instructed to provide their responses prior to reappearance and responses after reappearance were counted as misses. One-hundred and twenty out of 400 trials contained the incongruent displacement time and the number of trials were counterbalanced across conditions.

### 2.3 fMRI Data Acquisition

The scanning sessions were conducted on 3 Tesla Siemens PRISMA MR-system (Siemens, Erlangen, Germany), using a 64-channel head coil. The data of participants were acquired in 16 functional runs divided in two sessions, totalizing 1904 volumes for the training phases and 3144 volumes for the test phases for each subject^1^. Blood oxygenation level-dependent (BOLD) signals were acquired using a multi-band accelerated T2*-weighted echo-planar imaging (EPI) sequence (multi-band acceleration factor 2, repetition time (TR)=2000 ms, echo time (TE)=30 ms, flip angle=80°, field of view (FoV)=220 mm, voxel size=2.2 × 2.2 × 2.2 mm, no gap). Volumes were acquired in interleaved order. Identical slice selection on both days was achieved using Head Scout Localizer whose calculation is based on Autoalign (Siemens, Erlangen).

A high-resolution three-dimensional T1-weighted anatomical map (TR = 2500 ms, TE = 2.82 ms, FoV = 256 mm, flip angle = 7°, voxel size = 1 × 1 × 1 mm, 192 slices, parallel imaging with a GRAPPA factor of 2, and 5:18 min scan duration) covering the whole brain was obtained using a magnetization-prepared rapid acquisition gradient echo (MPRAGE) sequence. This scan was used as anatomical reference for the EPI data during the registration procedure.

### 2.4. Retinotopic Mapping

The procedure used for measuring the retinotopic maps was the similar to the one used by Warnking et al. (2002) and Bordier et al. (2015). Eccentricity was mapped using a checkerboard ring which slowly contracted or expanded from the fixation dot, while presented on a grey background. The speed of the expansion and the contraction varied linearly with the eccentricity, so that the activation wave kept travelling at an approximate constant speed (Bordier et al., 2015). The ring reached a maximum diameter eccentricity of 6.6 deg and a minimum of 0.2 deg. When the maximum (expansion) or the minimum (contraction) was reached, a new ring would start from the origin. Polarity was mapped using one checkerboard wedge (10 deg) slowly rotating at a constant speed. Specific parameter calculations were similar as the ones described by Warnking and colleagues (2002). The checkerboard stimulation flickered at a frequency of 8 Hz, in 10 cycles of 36 s each. The aspect ratio of the checkboards was kept constant (1.09) by scaling the height linearly with the eccentricity. In order to account for the effects of the hemodynamic delay, the wedges were presented clock- and counter-clockwise, and the rings were presented expanding annuli and contracting annuli (Warnking et al., 2002). In total, eight functional runs were acquired, two for each modality and direction, and each run lasted approximately 6 minutes. Data acquisition was done using a multi-band accelerated T2*-weighted EPI sequence (multi-band acceleration factor 2, TR=2000 ms, TE=30 ms, flip angle=90°, FoV=128 mm, voxel size=2.2 × 2.2 × 2.2 mm, no gap). We acquired 180 volumes in interleaved order for each run.

## 3. STATISTICAL ANALYSIS

### 3.1. Behaviour

Participants’ performance was assessed through the averaged correct responses (accuracy), response time and response time error (difference between individual’s response time and the presented stimulus duration). Trials with RTs exceeding 0.6s were excluded. These trials may have been contaminated by the information from the re-appearing stimulus and had a mean RT=0.864, (SE)± 0.027). On average 7.14 (SD± 9.83) of trials were excluded per subject.

The three measurements were calculated for training and test phases and used as input for three different repeated measures (rm) ANOVA: (1) for HP context, we used a 2×2 within-subject design (direction: up-down vs. velocity: fast-slow); (2) for LP context, we used a 2×2×2 within-subject design (direction: up-down vs velocity: fast-slow vs. congruency: congruent-incongruent); (3) for the comparison of HP vs. LP contexts, we carried out a 2×2×2 within-subject design as well (direction: up-down vs. velocity: fast-slow vs. predictability: high vs. low predictable) using the congruent trials only, as no incongruent trials were present in the HP context. Note that, when presenting the LP context results, we refer to the trials as incongruent fast whenever the stimulus travelled the horizontal trajectory fast and was *slowed-down* during the vertical trajectory in the visible or occlusion phases, and as incongruent slow when the stimulus travelled the horizontal trajectory slow and was *speeded-up* during the vertical trajectory in both phases, i.e. we use the horizontal speed for our terminology. Additionally, for each analysis, we included task order as a between subject factor, however in no statistical analysis any significant effect was observed (all p’s > 0.213). All analyses were calculated using JASP (v.0.15.0 - https://jasp-stats.org/). JASP was also used to compute post hoc tests and effect sizes (partial ƞ^2^).

### 3.2. Retinotopy

We performed a three-dimensional reconstruction of the cortical sheet based on the structural image of each of the 16 subjects using recon-all function from Freesurfer (v.6 - https://surfer.nmr.mgh.harvard.edu/). Retinotopic maps along the polar and eccentricity dimensions were calculated for each of the cortical surfaces using the “selxavg3-sess” function from Freesurfer. Lower and upper primary visual area were delineated manually on the flattened cortical sheets based on the boundaries of phase reversals within the polar angle and eccentricity maps (Abdollahi et al. 2014). Delineation of borders were created based on of Georgieva et al. (2009) and Kolster et al. (2010). The regions were later used to identify the local maxima during the visible phase in HP context. These local maxima were thus used to independently localise the region of interest within the functional region for the occluded phase. Probabilistic maps of MT as provided by Freesurfer parcellation was used for the identification of motion sensitive regions.

### 3.3. fMRI preprocessing

All data (except retinotopic data) were analysed using SPM12 (www.fil.ion.ucl.ac.uk/spm, Wellcome Trust Centre for Neuroimaging, London, UK). The first five volumes of each run were discarded to allow for steady state magnetization. We performed slice-timing corrected and realignment (registered to the mean image) of all remaining functional volumes. Head motion parameters were later used as nuisance regressors in the general linear model (GLM). Finally, the structural image was coregistered (estimate and reslice) to first functional image of the first run. Resliced images were smoothed with a gaussian kernel of 6 mm.

### 3.4. fMRI data Modelling

Data of individual contexts (HP and LP) were modelled with a general linear model (GLM, Friston et al., 1995), which included the run-wise condition parameters, derivatives, and six motion regressors as nuisance covariates. In particular, regressors of each condition (up-fast, down-slow, up-slow, down-fast or up-slow, down-fast, up-fast, down-slow, and respective incongruent conditions of LP contexts) were modelled with the canonical hemodynamic response function (HRF), using the onset of the initial stimulus trajectory of each trial. Temporal and dispersion derivatives of each regressor were added to the model in order to account for variability in the onset response and shape (Friston et al., 1998). Estimated beta weights of the HRF of each participant were extracted using MarsBar 0.44 (Brett et al., 2002) from subject-specific lower and upper V1 masks (see below for details of retinotopic analysis). Importantly, voxels containing response signal related to the reappearance of the stimulus were excluded from the data. We performed three rmANOVAs: (1) for HP context, we used a 2×2×2 (direction, velocity, V1 Quadrant), (2) for LP context for, we used a 2×2×2×2 direction, velocity, V1 Quadrant and congruency) and for combined congruent HP and LP, a 2×2×2×2 (direction, velocity, V1 Quadrant and predictability). Task order was included in the analysis as an between-subject effect, but no significant effects for this factor were observed (all p’s > 0.078). A full rmANOVA (direction, velocity, predictability, V1 quadrant, task order) was run for comparisons in which all tasks were included, however no difference between tasks were observed. Results are presented in the appendix B, table 1. For the MVPA, we modelled single trial GLMs (Least Square Separate a (LSS) approach, Mumford, 2012 – script adapted from https://github.com/ritcheym/fmri_misc/blob/master/generate_spm_singletrial.m).

**Table 1.**
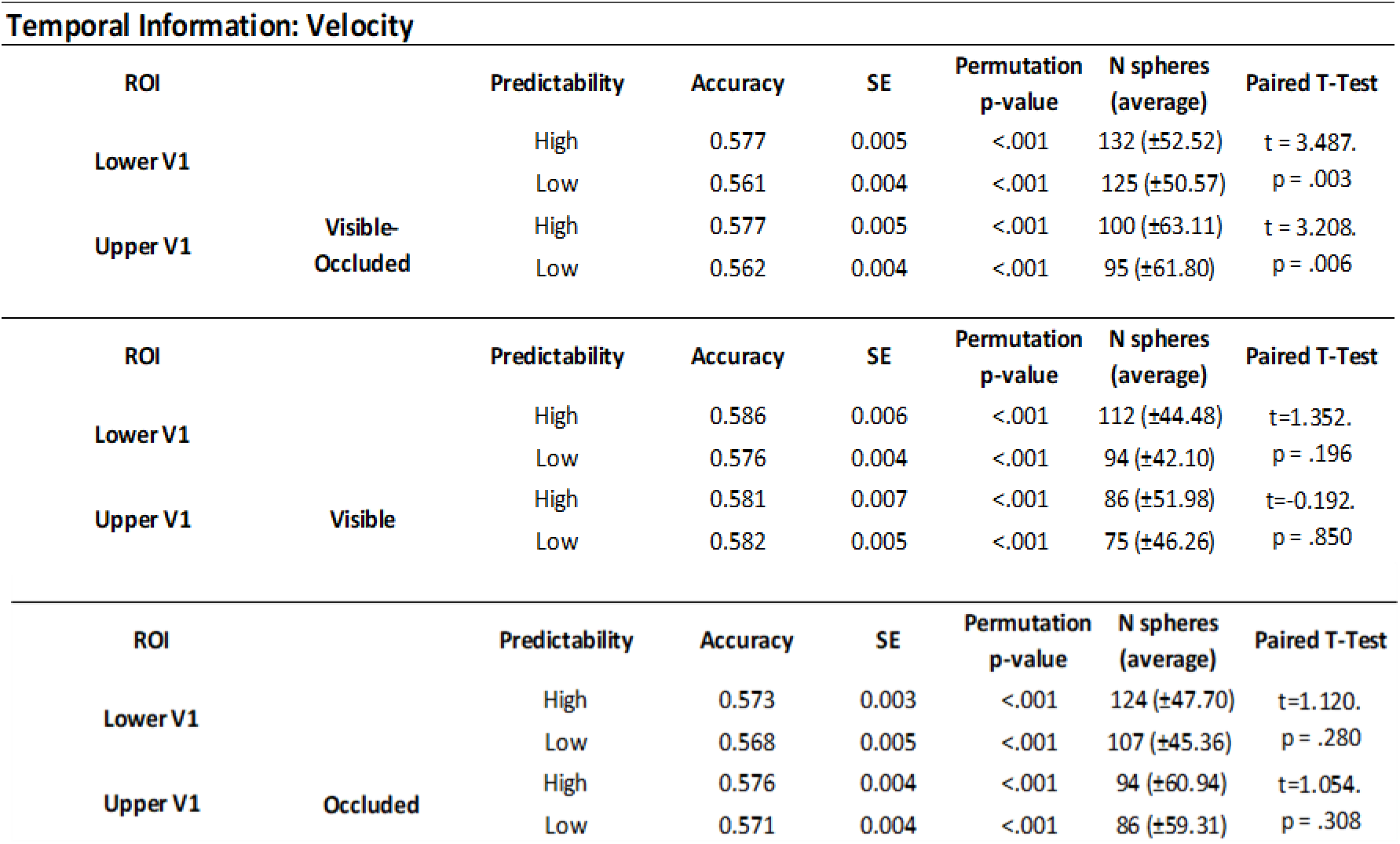
Decoding accuracy of temporal information: Accuracy values of all analyses of all conditions are displayed on the table above, together with the standard error of the mean (SE). Permutation p-values demonstrate that the permuted accuracy distributions were significantly lower than the true label accuracy distribution. The average number of spheres included in the sample of 5% most informative spheres can be seen in the range of 81 to 132.

### 3.5. Multivariate Pattern Analysis

A series of trial-wise multivariate pattern analyses was performed on beta values from GLMs of low and high predictable context for both Visible and Occluded Phases, using CoSMoMVPA (Oosterhof et al, 2016). To this end, trial-wise GLMs were carried out, and for the MVPA, the trials were calculated using Least Square Separate approach (Mumford, et al., 2012). Trial-wise MVPA was chosen here, due to the low number of runs for each task. Two runs are not enough to make valid train and test partitions as we would have only one in each part. For these cases, trial-wise analyses are recommended as the number of trials allow enough data in each partition (Mumford, et al., 2012). For all analyses, we used the searchlight method with a 4.4 mm sphere, a linear discriminant analysis (LDA) classifier and leave-25%trials-out (Etzel & Braver, 2013). All partitions were balanced and repeated 4 times. All analyses were carried out at the single-subject level, as the searchlight analyses were performed inside each individual mask.

We applied a cut-off at the accuracy of 0.5 (chance level) to filter out all the voxels which contained below chance accuracies, and included in the analysis only the values which belonged to the highest 5% values of the distribution (Agostino et al, 2023). This additional thresholding was done in order to obtain only the most informative voxels of the decoding. This procedure was applied to all the classification analyses below described. Permutations were carried out at the subject-level and 1000 iterations which contained randomised data labels per run, keeping the same original dataset. In order to have the spatial comparison, the same searchlight spheres included in the 5% highest accuracy sample were obtained for all 1000 samples of each individual participant. These sampled values were averaged across spheres for the original and permuted dataset permutation. For group level analysis, analyses were done based on Etzel (2017) approach. The null distribution carried the average across participants for each of the 1000 permutations plus the true-labelled group-level average (1001 group-level accuracies). The permutation p value was computed by taking the sum of the permuted accuracies higher or equal to the true-labelled accuracy and dividing by the number of iterations plus 1. Each analysis described below served a different purpose, such as decoding spatial, temporal information from both contexts, and congruency information from low predictable context.

#### Classifying Direction in HP & LP Context

Here, we carried out two classification analysis in which the classifier was trained in the visible phase and tested in the occluded phase using data of each context independently. We attempted to decode spatial visible information - upward vs downward motion trajectory - from occluded phase and compared accuracies from both analyses. We expected accuracies from LP context classification analysis to be significantly lower than HP contexts, due to the presence of incongruent trials, i.e. unpredictability.

#### Classifying Velocity in HP & LP Context

As the classification analyses above, we trained the classifier on the data of the visible phase and tested on data of the occluded phase within contexts. In this case, we attempted to decode temporal visible information - fast vs slow - from occluded data and compared accuracies from the different contexts. Here, we also expected decoding accuracies from LP context to be smaller than accuracies from HP context, also due to the presence of incongruent trials.

#### Manipulation checks

To verify whether the classifier was really decoding relevant information, classification analyses were performed on the visible and occluded data separately for both contexts. For HP contexts, direction and velocity were classified from both phases, as well as for LP contexts. Additionally, for the latter, congruent vs. incongruent information were classified. Here, we expected that decoding accuracies from visible phases were higher than occluded phases, as the longer exposure of the stimulus during the visible phase may allow a more robust representation of the information, whereas during occlusion, participants are expected to mentally represent the trajectory, thus less bottom-up input would drive the response.

We further ran pairwise Student’s T-test to compare HP and LP contexts in all conditions in lower and upper V1 quadrants to investigate whether conditions in LP context would encode more or less information, hence higher or lower accuracy values, compared to HP context.

## 4. RESULTS

### 4.1. Behaviour: Temporal Estimation

#### Visible Phase

##### High predictable context

Statistical analysis revealed main effects of velocity (F(1,14)=78.964, p<.001, ŋ_p_=.849; fast vs. slow: MD=-0.112, ±0.013, t=-8.886, p^bonf^ <.001) and direction (F(1,14)=15.200, p =.002, ŋ ^2^ =.521; upward vs. downward: MD=-0.023, ±.006, t=-3.889, p_bonf_ =.002), indicating that participants answered slower to slow motion, as expected, and also to the downward direction. Figure 2A shows averaged reaction time for all conditions during visible phase, for completeness.

**Figure 2.**
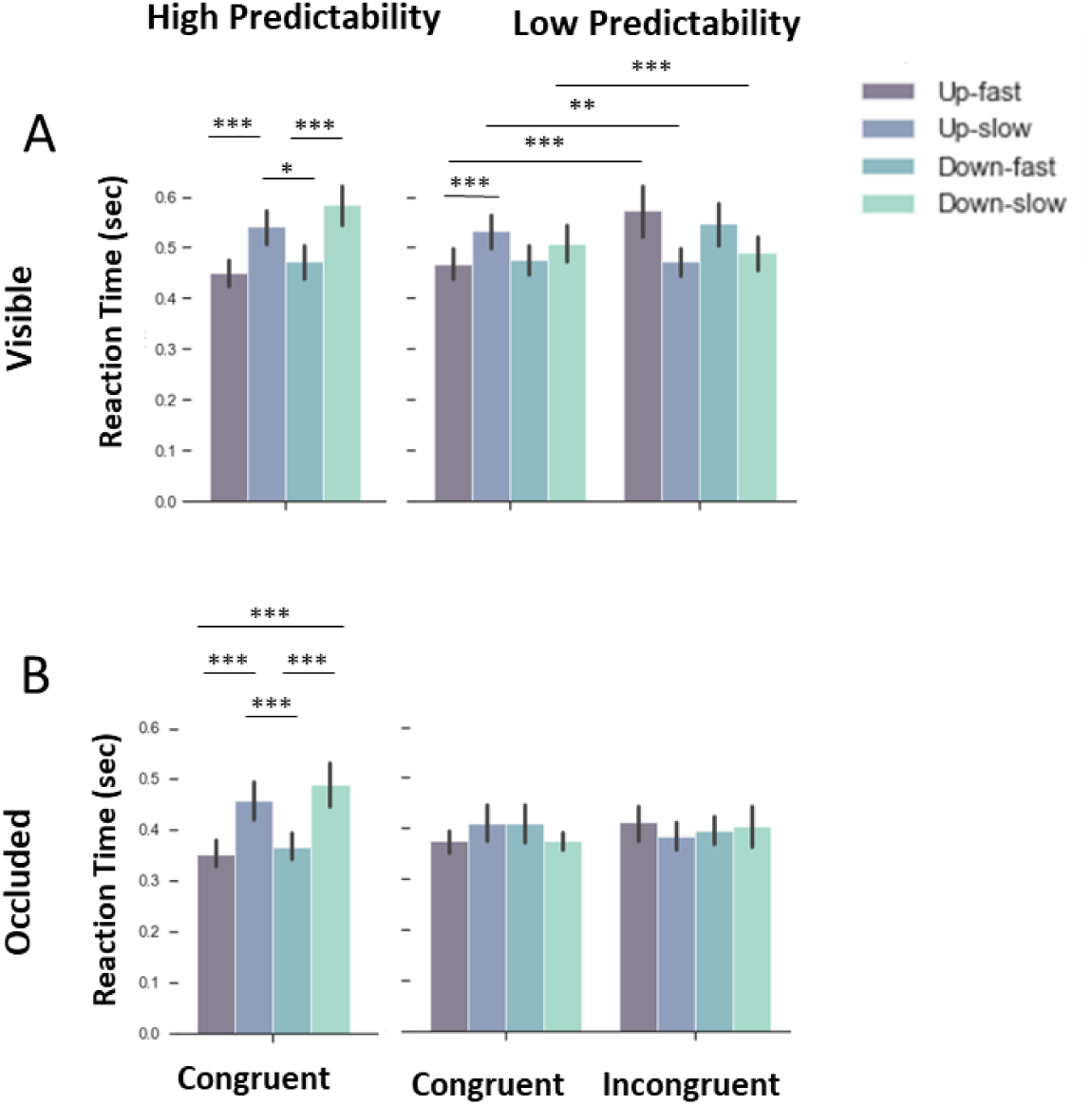
Group mean reaction times. Dark purple bars depict results of fast motion along the upward trajectory, blue bars depict results of slow motion along the upward trajectory, dark green bar, fast motion along the downward trajectory and light green, slow motion along the downward trajectory. (A) Reaction time results of the visible period in HP (left) and LP contexts (right). We observed differences between fast and slow conditions along the up- and downward trajectories, indicating that participants were estimating the time-to-contact correctly (answering faster when the stimulus was fast and answering slower when the stimulus was slow). This pattern can be clearly observed during in HP and LP contexts. (B) Results of the occluded period in HP (left) and LP contexts (right). Reaction times in the occluded HP context mirror those of the visible HP context, whereas this similarity was not found for the occluded LP context (B, right side).

##### Low predictable context

Figure 2A (LP context - Congruent) shows a similar RT pattern for the congruent LP context compared to the HP context. During incongruent trials the pattern apparently reversed, with stimulus slowing down in the vertical trajectory, showing even longer RTs than the slow trials in the congruent condition plus an effect of direction (Fig. 2A - LP context - Incongruent). Accordingly, the statistical analysis revealed a triple interaction between direction, velocity and congruency, which indicated slower response time during incongruent in both directions, and for congruent slow in upward direction (F(1,14)=5.882, p=.029, ŋ_p_^2^=.296; congruent fast vs. incongruent fast in upward: MD=-0.106, ±.014, t=-7.539, p_bonf_<.001; congruent fast vs. incongruent fast in downward: MD=-0.073, ±.014, t=-5.168, p_bonf_<.001; congruent slow vs. incongruent slow in upward: MD=.061, ±.014, t=4.313, p_bonf_=.002; see supplementary information for further results).

##### High x Low Predictability

While the pattern of results appears to be qualitatively similar for HP and LP congruent trials, the reaction times in the LP context appear to be compressed, with slow stimulus being faster and fast stimulus being slower in the vertical trajectory. Accordingly, we observed a triple interaction between direction, velocity and predictability (F(1,14)=5.340, p=.037, ŋ_p_^2^ =.276, see supplementary information for further results).

#### Occluded Phase

##### High Predictable Context

RT Results indicated main effects of velocity (F(1,14)= 78.964, p<.001, ŋ_p_^2^ =.849; fast vs. slow: MD=-0.112, ±.013. t=-8.886, p_bonf_ <.001), as well as of direction (F(1,14)= 15.200, p= .002, ŋ_p_^2^ =.521; upward vs. downward: MD=-0.023, ±.006. t=-3.899, p_bonf_ =.002), pointing to higher response time to slow motion and downwards direction, similar to the pattern observed during HP visible phase. Figure 2B shows averaged reaction time for all direction-velocity paired conditions during occluded phase.

##### Low Predictable Context & High x Low Predictability

Our statistical analysis did not yield any significant results for reaction time during low predictable context, suggesting that participants used an average of the slow and fast condition. A similar tendency was already observed during the LP visual phase with the RT compression.

Accordingly, for the direct comparison of response times in LP and HP contexts we found an interaction for velocity and predictability (F(1,14)=83.224, p<.001, ŋ_p_^2^ =.856; fast vs. slow in HP context: MD=-0.112, ±.010, t=-10.817, p_bonf_<.001; slow in HP vs. slow in LP context: MD=.079, ±.013, t=6.015, p_bonf_ <.001), suggesting higher response time for estimation for slow velocity during HP context, but lower response times for fast velocity in the HP context. This suggests that participants may have chosen a different strategy, as response times for the congruent trials showed a central tendency (see supplementary material for further details also of the Temporal estimation error and Spatial estimation which were always close to ceiling).

### 4.2. Univariate fMRI-results

#### Visible Phase

##### High predictable context

Enhanced fMRI signals along the upward trajectory were found in lower V1 and vice versa, in accord with established theories of visual processing. Accordingly, a statistical analysis of beta weights during visible stimulation presented in the HP context indicated an interaction between direction and V1 Quadrant (Fig. 3A), which suggests that the vertical stimulations were salient enough to elicit robust responses in the opposite quadrant (F(1,14)=76.090, p<.001, ŋ_p_^2^=.845; upward vs. downward in the upper V1: MD=-2.485, ±.511, t=-4.865, p_bonf_ <.001; upward vs. downward in lower V1: MD=2.230, ±.511, t=4.364, p_bonf_ <.001). Moreover, a main effect of velocity was found (F(1,14)=35.717, p<.001, ŋ_p_^2^=.718; fast vs. slow: MD=3.690, ±.617, t=5.976, p_bonf_<.001), as well as a triple interaction between direction, velocity and V1 quadrant (Fig. 4A), mainly pointing to higher fMRI-responses during fast downward motion in upper V1 (F(1,14)=6.761, p=.021, ŋ_p_^2^=.326; downward fast vs. downward slow in upper V1: MD=5.019, ±1.442, t=3.480, p_bonf_ =.043; downward fast in upper V1 vs. downward fast in lower V1: MD=4.200, ±1.155, t=3.636, p_bonf_ =.031, upward slow vs. upward fast in lower V1: MD=5.252, ±1.442, t=3.642, p_bonf_ =.028). No effect for task order (i.e. participant starts with HP or LP context) was observed for any of the conditions.

**Figure 3.**
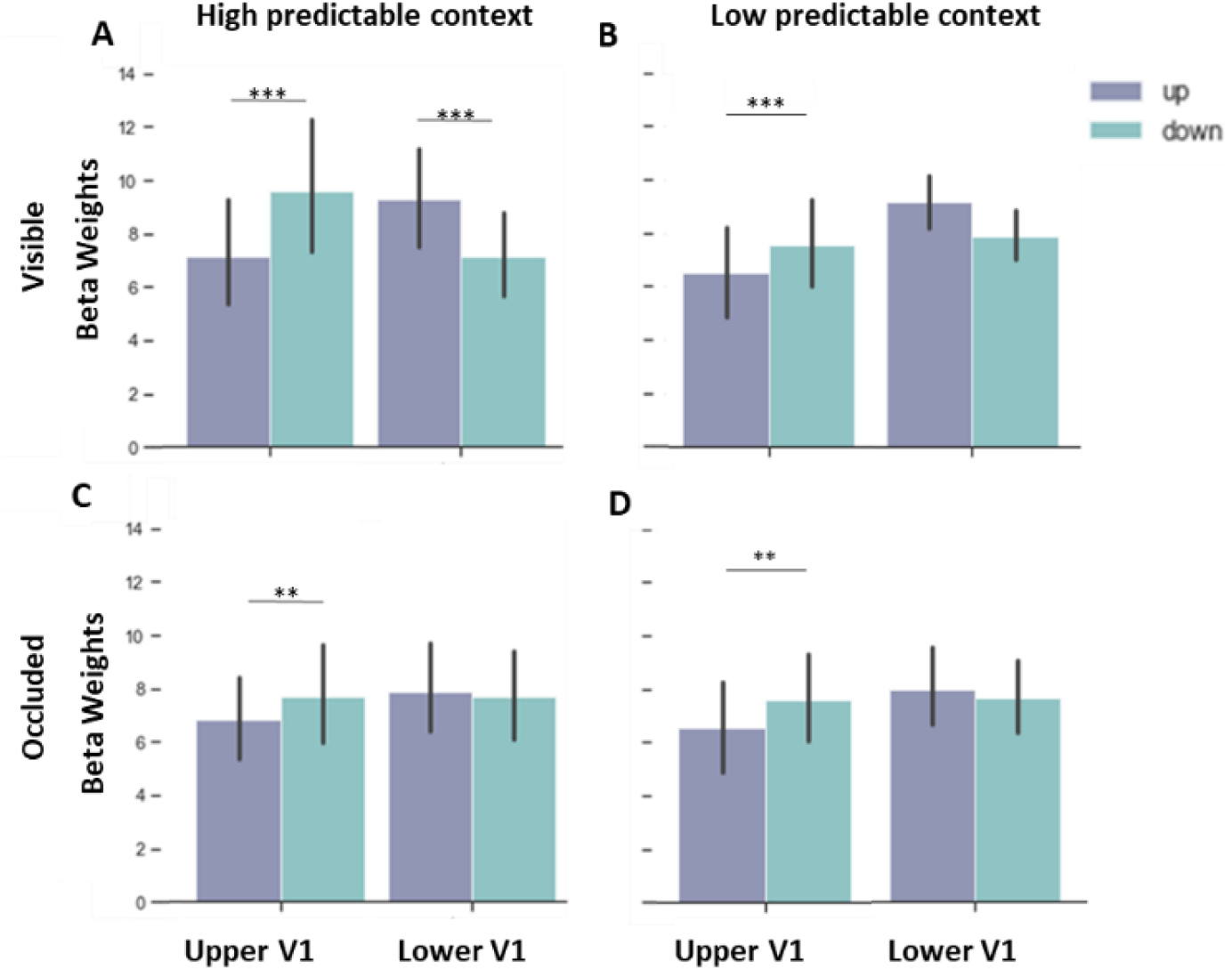
Effect of direction on V1-quadrant specific BOLD-responses. Beta weights (proportional to % signal change) of HP and LP contexts during (A) visible and (B) occluded phase, indicating an interaction between direction and V1 Quadrant. Purple bars represent upward trajectory and green bars represent downward trajectory. (A) During the visible phase in HP context, as expected, we observed that downward trajectory enhanced responses in upper V1, compared to upward trajectory, whereas upward trajectory elicited higher responses in lower V1 compared to downward trajectory. (B) During the visible phase in LP context, we observed the same pattern as in HP context, however, significant results were seen only in upper V1, while results in lower V1 were just marginally significant.

**Figure 4.**
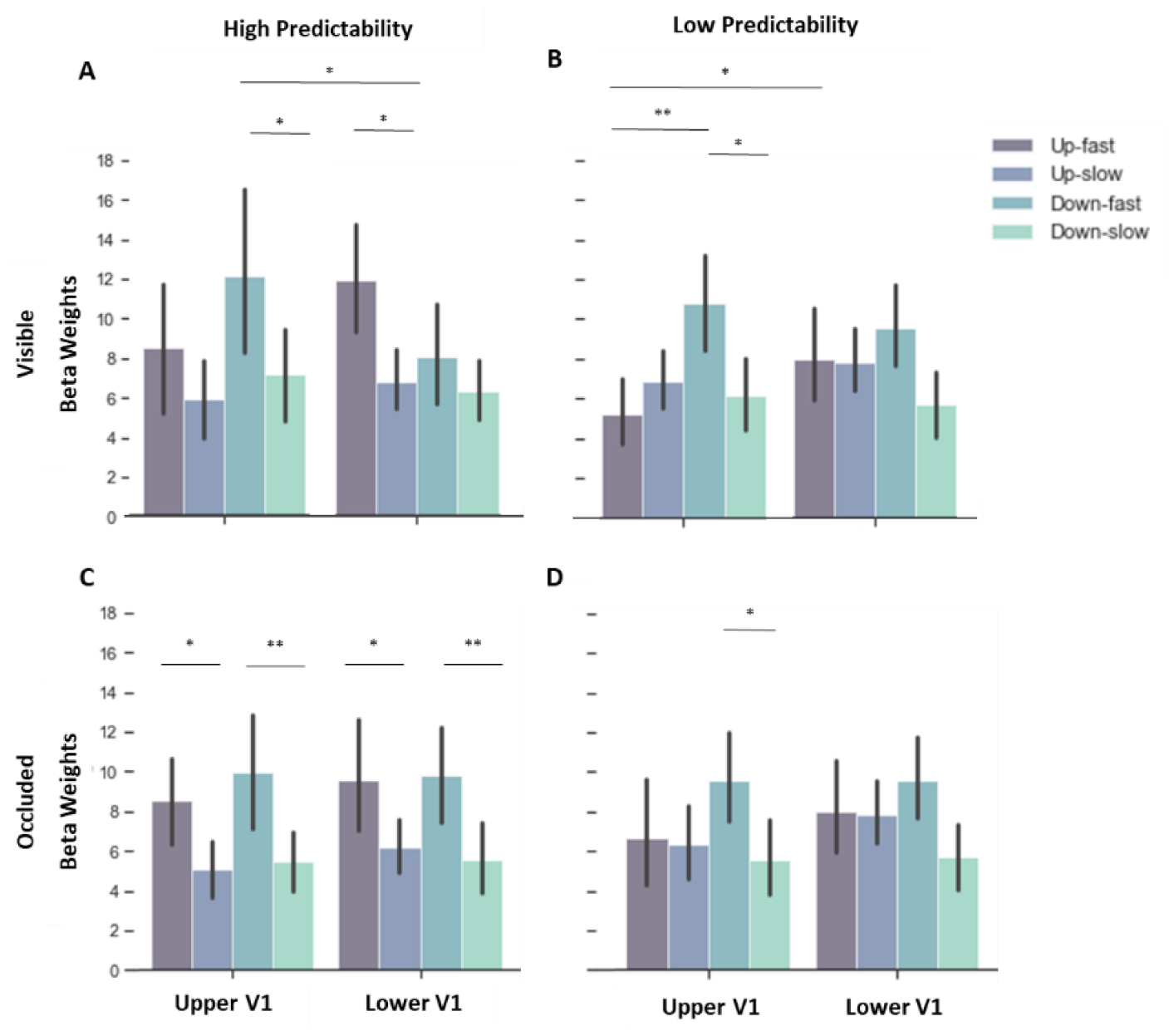
Beta weights (proportional to % signal change) of HP and LP contexts during the visible and occluded phase, indicating an interaction between direction, velocity and V1 Quadrant. Purple bars represent upward trajectories and green bars represent downward trajectories. (A) During the visible phase in HP context, we observed higher responses for fast compared to slow motion along downward trajectories in upper V1, which were also higher compared to fast downward in lower V1. In lower V1, responses during fast motion were higher than slow motion along upward trajectories. (B) During visible phase in LP context, fast motion compared to slow motion elicited higher responses for downward trajectories in upper V1, which were also higher compared to fast motion in upward direction. However, fast upward motion was higher in lower V1 compared to upper V1. (C) During the occlusion phase in HP context, differences were observed within the same quadrant only. Fast motion in upward, as well as in downward trajectory elicited higher responses compared to slow motion for the respective trajectories. (D) In LP context, during occlusion phase, fast motion compared to slow motion in downward direction enhanced responses in upper V1.

##### Low predictable context

During visible stimulation in the low predictable context, we observed again direction and V1 Quadrant interaction (Fig. 3B - F(1,14)=26.231, p<.001, ŋ_p_^2^=.652), which demonstrated that, the insertion of the noise through the incongruent trials still led to higher responses in the opposite quadrant; this effect was significant for upper V1 (MD=-2.630, ±.472, t=-5.573, p_bonf_ <.001), and marginally significant for lower V1 (upward vs. downward in lower V1: MD=1.304, ±.472, t=2.764, p_bonf_ =.063). Indeed, in the congruent condition fast motion elicited higher responses in V1 than slow motion, whereas the incongruent trials with changes in velocity led to intermediate responses compared to fast and slow congruent trials. Statistically, an interaction between velocity and congruency (Fig. 5A) was found with higher fMRI-signals during congruent fast motion and incongruent slow motion (F(1,14)=99.931, p<.001, ŋ_p_^2^=.877; congruent fast vs. slow: MD=4.240, ±.374, t=11.329, p_bonf_ <.001; congruent fast vs. incongruent fast: MD=3.319, ±.455, t=7.286, p<.001; congruent slow vs. incongruent slow: MD=-1.706, ±.455, t=-3.746, p_bonf_ =.006).

**Figure 5.**
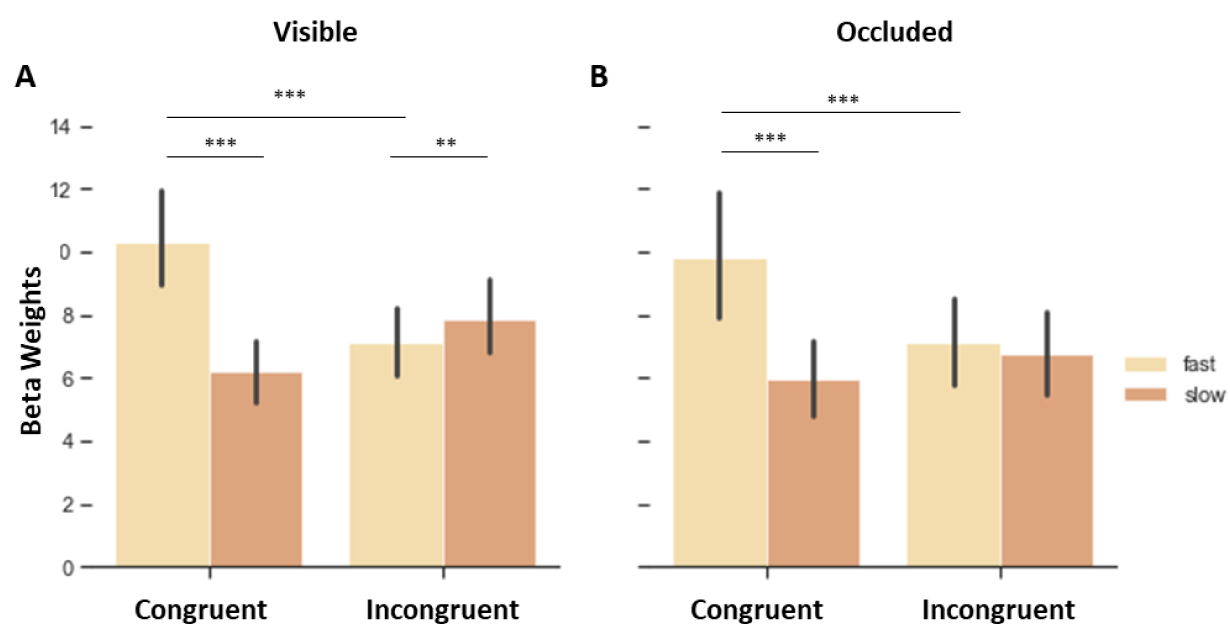
Beta weights of congruent and incongruent trials in visible and occlusion phases. Fast congruent trials elicited enhanced fMRI-responses in V1 during (A) visible and (B) occluded phases, compared to slow congruent and incongruent conditions. Note, however, that slow incongruent (speeded-up stimulus in the vertical trajectory) elicited higher responses compared to fast incongruent (slowed-down stimulus in the vertical trajectory).

We further observed an interaction between direction and velocity (F(1,14)=4.694, p=.048, ŋ_p_^2^=.251; downward fast vs. downward slow: MD=4.212, ±1.180, t=3.570, p_bonf_ =.016), and a marginally significant triple interaction between direction, velocity and V1 Quadrant (Fig. 4B), which yielded significant post-hoc comparisons (F(1,14)=4.240, p=.059, ŋ_p_^2^=.232; upward fast vs. downward fast in upper V1: MD=-5.799, ±1.284, t=-4.517, p_bonf_ =.005; downward fast vs. downward slow in upper V1: MD=4.766, ±1.237, t=3.852, p=.031).

##### High x Low Predictability

When comparing beta weights of congruent trials from both contexts, we observe interactions between direction and V1 Quadrant (F(1,14)=54.439, p<.001, ŋ_p_^2^=.795; upward vs. downward in the upper V1: MD=-2.777, ±.456, t=-6.087, p_bonf_ <.001; upward vs. downward in lower V1: MD=1.515, ±.456, t=3.320, p_bonf_ =.016), which indicated that the vertical stimulations were salient enough to elicit higher responses in the opposite quadrant. Additionally, triple interactions were seen between direction, velocity and V1 Quadrant again highlighting that velocity played a role in V1 responses (F(1,14)=12.864, p=.003, ŋ_p_^2^=.479; upward fast vs. upward slow in lower V1: MD=4.075, ±1.137, t=3.585, p_bonf_ =.004; upward fast vs. downward fast in upper V1: MD=-4.965, ±1.118, t=-4.441, p_bonf_=.005; downward fast vs. downward slow upper V1: MD=6.158, ±1.137, t=5.418, p_bonf_<.001) and velocity, V1 quadrant and predictability (F(1,14)=6.408, p=.024, ŋ_p_^2^=.314; though no significant effects were found in the post-hoc comparisons between different contexts). Finally, a main effect of velocity was observed (F(1,14)=78.946, p<.001, ŋ_p_^2^=.849; fast vs. slow = MD=3.965, ±.446, t=8.885, p_bonf_ <.001).

#### Occluded Phase

##### High predictable context

Beta weights of the high predictable occluded phase indicated an interaction between direction and V1 Quadrant (Fig. 3C), indicating that during occlusion, downward direction elicited a similar pattern of responses in upper V1, but not lower V1, compared to the visible phase (F(1,14)=10.927, p=.005, ŋ_p_^2^=.438; upward vs. downward in upper V1: MD=-0.945, ±.285., t=-3.315, p_bonf_=.016). Further, a main effect of velocity was found (F(1,14)=42.861, p<.001, ŋ_p_^2^=.754; fast vs. slow: MD=4.013, ±.613, t=6.547, p<.001). Since we observed the triple interaction between direction, velocity and VF in the visible phase (Fig. 4C), we also computed T-tests for the occluded phase and found significant differences between upward fast vs. upward slow in lower V1 (MD=4.013, ±.613, t=6.547, p=.044); downward fast vs. downward slow in lower V1 (MD=4.230, ±1.054, t=4.015, p=.009); upward fast vs. upward slow in upper V1 (MD=3.667, ±.1.054, t=3.480, p=.039); downward fast vs. downward slow in upper V1 (MD=4.531, ±1.054, t=4.301, p=.004).

##### Low predictable context

During the low predictable context, we once more observed an interaction between direction and V1 quadrant (Fig. 3D - F(1,14)=19.683, p<.001, ŋ_p_^2^=.584; post-hoc comparisons revealed the difference only when incongruent trials were not included in the analysis: F(1,14)=14.691, p=.002, ŋ_p_^2^=.517; upward vs downward in the upper V1: MD=-1.532, ±.436, t=-3.511, p=.014). Additionally, interactions between velocity and congruency were found(Fig. 5B - F(1,14)=30.281, p<.001, ŋ_p_^2^=.684; congruent fast vs. congruent slow: MD=3.953, ±.602, t=6.561, p_bonf_<.001; congruent vs. incongruent fast: MD=2.767, ±.461, p_bonf_<.001), as well as direction and congruency (F(1,14)=15.575, p=.001, ŋ_p_^2^=.527; congruent vs. incongruent downward: MD=1.462, ±.349, t=4.192, p_bonf_=.003). Further, results revealed a marginally significant interaction between direction and velocity (F(1,14)=4.016, p.065, ŋ_p_^2^=.223; downward fast vs. slow: MD=4.119, ±1.101, t=3.743, p_bonf_=.007. For the sake of completeness, we computed post-hocs t-Tests on the triple interaction between direction, velocity and VF (Fig. 4D) and observed significant difference between downward fast vs. downward slow in upper V1 (MD=4.288, ±1.171, t=3.610, p_bonf_=.035).

##### High x Low Predictability

During occlusion, the comparison between both contexts yielded significant interactions of direction and V1 quadrant (F(1,14)=18.134, p<.001, ŋ_p_^2^=.564; upward vs. downward in the upper V1: MD=-1.238, ±.301, t=-4.116, p_bonf_=.003), and a marginally significant interaction of direction and velocity (F(1,14)=3.451, p=.084, ŋ_p_^2^=.198; upward fast vs. upward slow: MD=2.701, ±.894, t=3.020, p_bonf_=.033; downward fast vs downward slow: MD=5.265, ±.894, t=5.889, p_bonf_<.001). We further observed a main effect of direction (F(1,14)=5.489, p=.034, ŋ_p_^2^=.282; upward vs. downward: MD=-0.616, ±.263, t=-2.343, p_bonf_=.034) and velocity (F(1,14)=49.106, p<.001, ŋ_p_^2^=.778; fast vs. slow: MD=3.983, ±.568, t=7.008, p_bonf_<.001), but no main effect of or interaction with predictability.

### 4.3. Trial-by-trial Information in LP context

The results above showed that there are differences between congruent and incongruent trials in both visible and occluded conditions. However, differences in BOLD-response might not only be due to the cumulative trial-history (i.e. LP vs. HP context) but to the recent trial-history. To this end, we analysed the trial-by-trial fMRI-responses in V1 to test whether responses to congruent trials after incongruent trials (and vice-versa) are higher, due to lower predictability (i.e. congruent followed by congruent or incongruent followed by incongruent trials - Fig.6). We performed a repeated measures ANOVA (2×2×4: visible/occluded; V1up/V1down; congruence: congruent trials followed by incongruent trials [CI], incongruent trials followed by congruent trials [IC], congruent trials followed by congruent trials [CC], incongruent trials followed by incongruent trials [II]). We found significant effects only for congruence (F(3,45)=4.678, p=0.006, ŋ_p_^2^=.059; CI vs. CC: MD=-0.894, ±.260, t=-3.441, p_bonf_=.008; IC vs. CC: MD=-0.774, ±.260, t=-2.978, p_bonf_=.028), and no difference between visible and occluded phases (F(1,15)=1.643, p=0.219, ŋ_p_^2^=.023).

**Figure 6.**
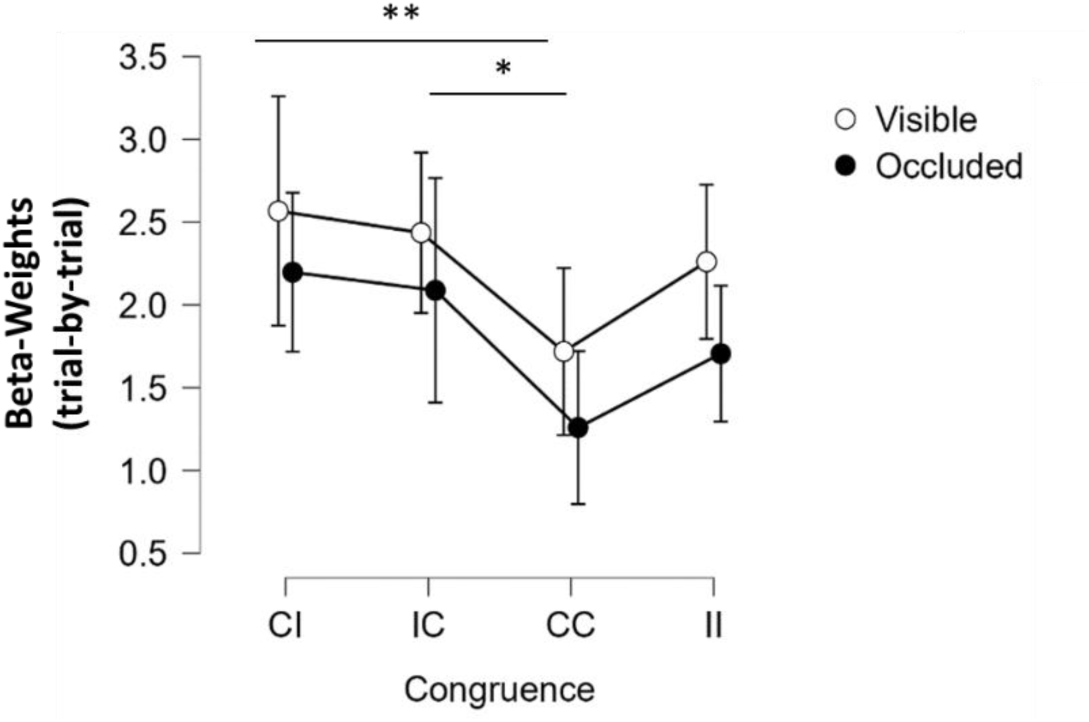
Trial-by-trial comparison for visible and occluded phases. The trial sequence abbreviations: CI = congruent trials after incongruent trials, IC = incongruent trials after congruent trials, CC = congruent trials after congruent trials, and II = incongruent trials after incongruent trials.

### 4.4. Multivariate Pattern Analysis results

To test whether the informational content in V1 was similar during visible and occluded conditions, we performed trial-wise searchlight MVPA analyses in HP and LP contexts separately. Additionally, a comparison between both contexts were carried out to directly compare the informational content in both contexts. A multivariate analysis of trial-history effects was not possible due to the low number of trials in some conditions. Table 1 and 2 depict the accuracy values of all analyses, of which we decoded spatial information, i.e. upward vs. downward trajectory; and temporal information, i.e. fast vs. slow motion, respectively.

#### 4.4.1. Temporal Information: classifying velocity

##### Classifying Velocity Patterns of occluded Phase from visual Phase in HP and LP contexts

Here, we trained the classifier in the visible phase and tested in the occluded phase data using fast and slow conditions as labels, separately for HP and LP contexts. Note that trajectory changes up/down were collapsed, so only velocity can possibly explain any results. We focused our analysis on the primary visual areas (see supplementary material for additional results in area V5/human MT+). Decoding accuracies, obtained from the sample of the 5% most informative spheres (range of 81 to 132 spheres) were above chance level (Tab. 1: Visible-Occluded), suggesting that temporal information is represented in visual areas. Importantly, the representational patterns of temporal information are similar during visible and occluded phases.

#### 4.4.2. Spatial Information: classifying direction

##### Classifying Direction Patterns of Visible Phase from Occluded Phase in HP and LP contexts

In these analyses we trained the classifier in the visible phase and tested in the occluded phase data using upward and downward conditions as labels, separately for HP and LP context. Note that the results from both sessions with opposing velocity association were collapsed, so that a decoding of stimulus velocity cannot explain the results below. Here we expected to decode information from the occluded data with training in the visible phase, as previously observed (Agostino et al., 2023). In order to obtain the most informative voxels, we averaged the 5% most informative spheres (range of 45 to 112 selected spheres). Accuracies above chance level were found, suggesting that we could successfully decode direction-specific informational patterns from the visible phase in the occluded phase in the lower and upper V1 and V5. Additionally, all analyses with the true labels yielded significantly higher accuracies compared to the permutation analyses (Tab. 2: Visible-Occluded). Together, these findings indicate that MVPA significantly extended the results from the univariate analyses by showing that both phases share similar informational pattern in the primary visual area. Furthermore, these classification analyses replicate the results of our previous study, in which we demonstrated a common engagement of low-level visual cortex during the presentation of visible and dynamically occluded motion.

**Table 2.**
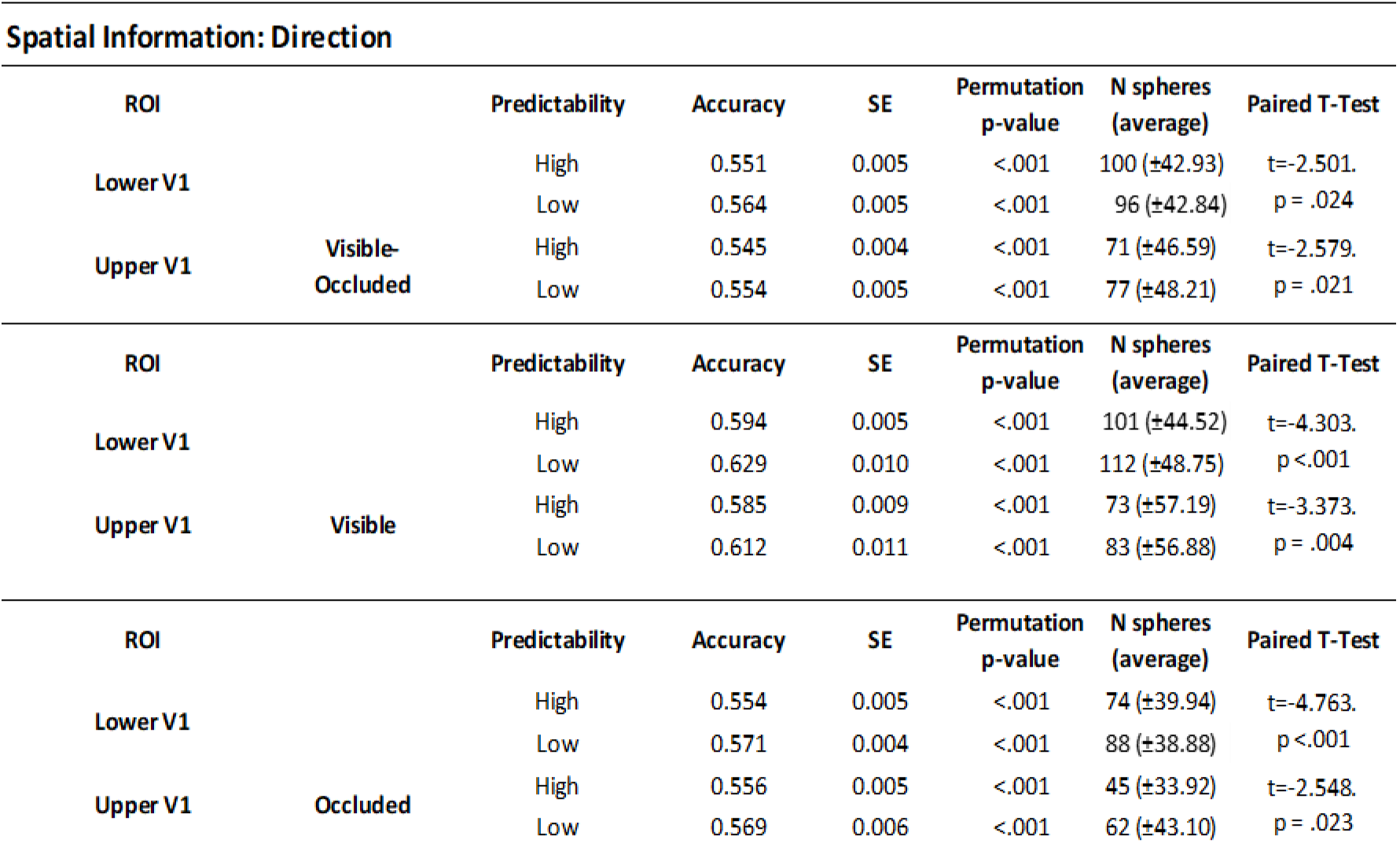
Decoding accuracy of spatial information: Accuracy values of all analyses of all conditions are displayed on the table above, together with the standard error of the mean (SE). Permutation p-values demonstrate that the permuted accuracy distributions were highly significant different from the true label accuracy distribution. The average number of spheres included in the sample of 5% most informative spheres were in the range of 45 to 112.

##### Manipulation checks

As a manipulation check for spatial decoding, we tested the classifier in visible and occlusion phases separately (Tab. 2: Visible; Occluded). Results indicated that accuracies from the classification analysis in which we trained and tested in visible phase data were higher compared to the two other classification analyses Such results were expected, as during the visible period more visual information is available to be encoded by the visual system. In contrast, the other two analyses presented very similar averaged decoding accuracies. Also note that the decoding accuracies of this analysis were generally smaller compared to a previous study (Agostino et al., 2023) as we used trial-wise rather than run-wise estimates as input to the MVPA, which includes an increased noise level, while the previous study used run-wise averages.

As manipulation check for the temporal decoding, we analysed visible and occluded phases separately. Note, however, that, in comparison with the decoding of spatial information, we did not observe higher accuracy for visible phase analysis compared to the other two analyses (Tab. 1: Visible; Occluded). This may indicate that the pattern of temporal information was encoded similarly, independent of the availability of the stimulus (visible or occluded). In contrast with the comparison of spatial information decoding accuracies from HP and LP context, in which we observed higher accuracies for the latter, accuracies from analyses with LP context were not higher than accuracies from analyses with HP context data (Table 2). Instead, classification analyses with visible phase only or occluded phase only did not yield significant different results between both contexts; whereas analyses in which the classifier was trained in visible and tested in occluded HP context data yielded higher accuracies than the same analysis with LP context data. These findings might suggest that temporal information can be better decoded when the encoding is not affected by some noise, such as the reliability of the stimulus velocity.

When comparing accuracies of HP and LP context, we observed significantly higher accuracies decoded from LP context data compared to accuracy decoded from HP context data in all analyses, this may suggest that the incongruent condition might have led to an encoding of more information. Therefore, we directly compared of congruent and incongruent decoding.

#### 4.4.3. Congruent Information in LP context

##### Classifying Incongruent trials of Visible Phase and Occluded Phase in LP context

These analyses were carried out by training the classifier in the visible and occluded data separately, using congruent and incongruent conditions as labels. Results (Fig.7) indicated that averaged accuracies (average of the number of the 5% most informative spheres is 75, ±41.05) were significantly above chance level compared to permutation analyses of visible phase in lower V1 (mean (M) = 0.572, ±.024, permutation p (p_perm_) <.001; upper V1 (M=.566, ±.019, p_perm_<.001) and V5 (M=.571, ±.022, p_perm_<.001), as well as of occluded phase in lower V1 (M=.554, ±.015, p_perm_<.001), upper V1 (M=.552, ±.014, p_perm_<.001) and V5 (M=.580, ±.023, p_perm_<.001). These results indicated that incongruent trials were successfully classified to be different from congruent trials, suggesting that different information was encoded during the tasks with lower temporal predictability. While this difference might be due to the observable changes in speed during the visible phase, this argument does not account for the differences found during the occluded phase. Finally, these findings significantly extend the univariate analysis which showed different pattern of activity between congruent and incongruent conditions.

**Figure 7.**
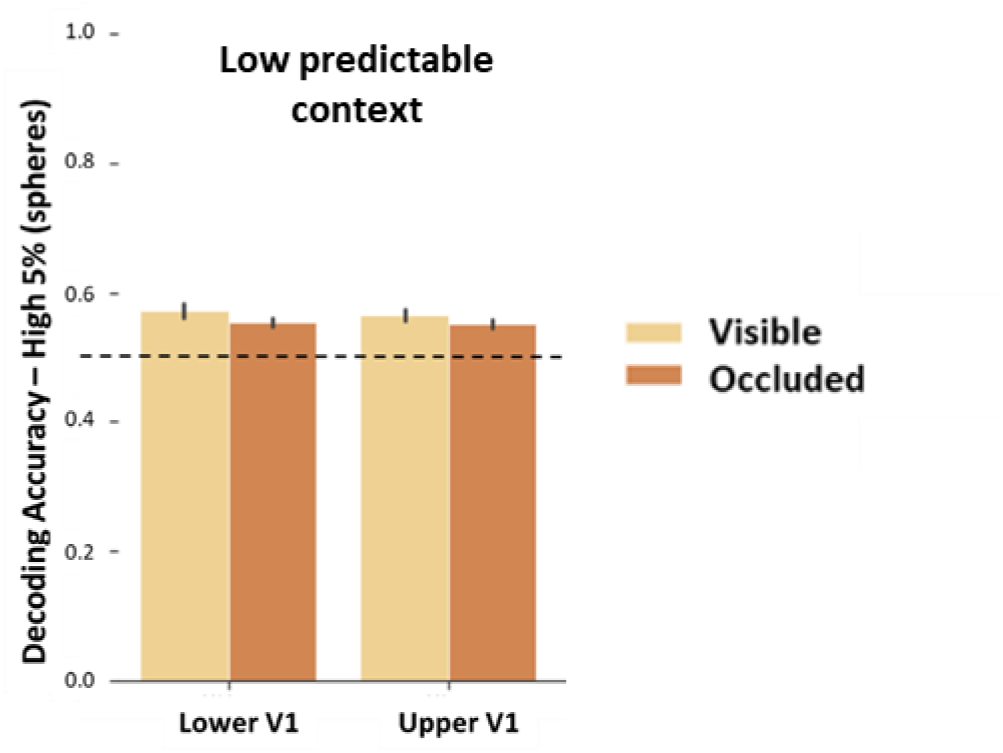
Classifying congruent and incongruent information. Yellow bars depict averaged accuracy of classification analyses in which the classifier was trained and tested in visible phase (“Visible”), orange bars depict the averaged accuracy of classification analyses in which the classifier was trained and tested in occluded phase (“Occluded”). The dashed line on 0.5 represent the theoretical chance level

#### 4.4.4. Overlap of sphere centres

In our previous study (Agostino et al., 2023) we used the 5% highest accuracy approach and showed that the overlap of significant spheres across analyses is above 80% and at the locations of V1 which represented the stimulus trajectory.

To evaluate that our approach of choosing the 5% highest accuracy values is robust and free of bias and does not randomly select different voxels within a region for the different analyses in he current study, we again calculated the number of spheres which share the same centres across different classification analyses. Interindividual results indicated that all participants shared spheres across different analysis in different probability context in all masks. The averaged overlapped spheres centre for all classification analyses performed here can be seen in tables 1 and 2. The overall average of all analyses is 83.19 (sd: ±21.63). Note that this value is similar to the number of sphere centres reported in the previous study (Agostino et al., 2023). This similarity between overall averages of both studies may be a result of the down-sampled space, as the analyses were carried out in the reduced space of ROI masks. Together, these results suggested that our approach of selecting the 5% highest informative voxels are robust, supporting the interpretation that when different decoding analyses share the same sphere centre, the decode pattern is truly informative, rather than a random selection of noise.

## 5. DISCUSSION

### 5.1. Behavioural analysis read-outs

This study investigated whether temporal motion extrapolation mechanisms would be affected by high vs low predictable contexts, and whether the behavioural effects are related to differential fMRI-responses in the primary visual cortex during visible and occlusion period. Our behavioural results demonstrated that during the visible phase, differences were observed between HP and LP contexts, indicating that performance was higher for HP compared with LP especially for slow motion in downward trajectory.

Reaction time results during visible stimulation showed that participants attempted to estimate the time-to-contact of the stimulus accordingly (faster responses during fast motion, slower responses during slow motion), in HP and LP context, following the expected pattern, and corroborating the results from our first study (Agostino et al., 2023). However, we, further, observed an interaction between velocity, predictability and direction which indicated that participants responded faster to slow downward motion during the LP context. During occlusion, temporal estimation in LP context did no longer follow the same pattern, meaning that faster trials were not estimated faster and slower trials were not estimated longer. Rather, reaction times were time compressed across conditions in LP context, suggesting that when accurate temporal estimation was impossible in 30% of the cases, participants used an averaging strategy. It could be speculated, that this effect might be mediated by post-error adaptations (Danielmeier & Ullsperger, 2011; King et al., 2010). Post-error adaptations alter future behaviour, leading to potential improvements, such as faster reaction times or higher accuracies. One of these adaptations is known as post-error speeding and it was showed that it is related to an enhancement of performance after a given threat (Caudek et at, 2015) and an increase of activity in task-relevant visual areas (King et al., 2010). However here, we didn’t observe the improvement of fast stimulus estimation, as participants were still trying to adequately respond according to the learned pattern. This was also found in a reversal of responses for incongruent relative to congruent trials during occlusion, pointed by reaction time error, which suggests that participants applied the learned congruent direction-velocity change (see supplementary material). Summing up, the pattern of behavioural temporal estimates indicate that participants performed well in both contexts in the visual phase with some overestimation of TTC in the fast condition in accord with previous publications. During occlusion, however, an increase in RT errors was observed for the LP context relative to the HP context.

### 5.2. Univariate analysis read-outs

At the neural level, we observed V1-quadrant-specific responses to upward and downward motion, as expected. No significant differences were found between HP and LP context, neither during visible nor during occlusion phase. This finding is not in line with our initial hypothesis of enhanced fMRI-signals in the LP context. However, previous studies which observed an enhancement in activity in the primary visual cortex during the processing of dynamically occluded stimulation due to attention did not modulate attentional demands parametrically (Kok et al., 2012; Coull et al. 2008; Doherty et al., 2005), thus it is conceivable that attention acts in an all-or-none way, and would therefore increase the signal to the maximum in both contexts. Hence, attention would affect prediction processing not gradually but categorically, as indicated by Fischer and colleagues (2013), who investigated the relation between temporal predictability and temporal attention and reported that V1 could be primed and more responsive to temporally predictable stimuli. Moreover, it is important to emphasize that during occlusion participants did not make a simple time-to-contact judgement, but rather they were required to make a more complex prediction due to the velocity-direction association. Potentially the reappearance of the stimulus may have automatically captured attention and thus may have generated a response high enough to decrease the pattern of activity related to the predictive error, thus equalizing the response pattern in HP and LP contexts. (see also below, where we discuss the results at the trial level which may support predictive coding o some extent).

We further observed that upwards trajectories enhanced responses in lower V1 and downward trajectories enhanced responses in upper V1, not only during the visible phase, but also and most importantly, during occlusion phase. These results corroborated our previous study (Agostino et al, 2023), which demonstrated that the mechanisms related to visualizing and extrapolating a moving stimulation indeed modulate activity in identical searchlights in low-level visual areas. However, here we observed that the expected pattern was seen in visible and occlusion phase in HP and LP contexts in upper V1, hence lower visual field. Differences between vertical hemifields have been reported previously. The vertical meridian asymmetry (Carrasco et al., 2001; Rijsdijk et al., 1980; Previc, 1990) indicates a dominance of the lower visual field over the upper visual field in different tasks, such as spatial resolution (Talgar & Carrasco, 2002), visual acuity (Skrandies, 1987), motion (Levine & McAnany, 2005), among others (Karim & Kojima, 2010, for review). An ecological explanation for this difference between vertical meridians was given by Previc (1990), who proposed that the dominance of the lower visual field comes from the primordial of the primate visual system, which was functionally more developed due to forelimb manipulatory skills. Later, studies indicated that the lower VF contains a larger amount of “near-preferring” neurons most commonly found in the latter compared to the first (Nasr & Tootell, 2018, 2020; Karim & Kojima, 2010). These findings could explain the difference between this and our previous study, given that in this study we introduced the reappearance of stimulus, which might also have led to an automatic capture of attention (Lakha & Humphreys, 2005). It is important to mention once more that the portion (voxels) in the respective quadrants representing the position of the reappearance of the stimulus were carefully excluded from the analyses to avoid misleading conclusions.

While in the spatial domain the stimulation modulated different patterns of activity, in the temporal domain univariate results were unanimous. The presentation of fast stimulation consistently enhanced activity in both upper and lower visual quadrants. Interestingly, even during incongruent conditions of visible stimulation, there was a tendency for the incongruent slow motion (speeded-up stimulus) to follow the same pattern of enhanced activity observed during the presentation of congruent fast motion. One possible explanation for these findings is that V1 may be more promptly receiving feedback projections from V5 during fast motion (Edwards et al. 2017; Sterzer et al., 2006; Muckli et al. 2005), which could increase the response signal, while processing of slow motion would be taking longer to reach V5 and reach V1 back. Note that the univariate results partially oppose to the behaviour results in which we observed that a major effect for slow, rather than fast motion. However, during LP context, the reversal pattern seen during incongruent trials in reaction time, could also be seen as a tendency in the univariate results.

Finally, our trial-by-trial analysis showed that beta values of trials following another trial of a different category (IC; CI) were higher than those of trials following trials of the same category (CC; II). These findings support our first hypothesis that unpredictable stimulus enhance activity in V1, in accord with the notion of the predictive coding theory, suggesting that the averaging of congruent and incongruent trials ameliorated the different representational patterns of the conditions in LP context. It is worth-mentioning that trial-by-trial GLM captures the temporal variability across trials, whereas the common GLM may average out some of these trial-specific effects (Mumford et al, 2012). Moreover, by modelling each trial separately, the hemodynamic response variability particular to each trial, and influenced by attention, arousal and task engagement, can be better captured compared to the common GLM, which may overlook or attenuate these trial-specific effects (Mumford et al, 2012; Rissman et al, 2004; Dale & Buckner, 1997). In sum, our univariate analyses revealed quadrant-specific effects in V1 during visible stimulation and to some extent during occlusion. Moreover, differential fMRI-responses were observed for the different velocities.

### 5.3. MVPA read-outs

The series of multivariate pattern analyses extended the results of the univariate analyses. First, we observed that representational patterns of activity from the visible phase could be used to decode patterns from the occluded phase in upper and lower V1, thereby extending the results from the univariate analyses, where we only found effects in upper V1. Our results suggest that visible and extrapolated types of information share similar representational pattern in the very same voxels, replicating the results from our previous study (Agostino et al. 2023). The appropriateness of the sphere selection approach that we used was further confirmed by computing the overlap of sphere centres. This overlap indicated that the majority of spheres shared the same centre in upper or lower V1 across different types of classification (train in visible, test in occluded; train and test in visible; and train and test in occluded), suggesting that the algorithm was decoding relevant information rather than noise, and also corroborating our previous study. Importantly, the effect was observed both in HP and LP contexts.

However, when comparing the accuracy values of HP with LP context, we observed significantly higher accuracy values in the LP context. These results may be due to enhanced attention in the LP context in which stimuli had to be monitored closely after they changed direction to detect a possible change in speed. Moreover, the change in velocity might have led to a difference in brain activity. Indeed, we observed a difference between congruent and incongruent trials, when we classified congruent and incongruent trials from visible and occluded data separately, confirming that the representational patterns of congruent and incongruent trials were significantly different. The results from the visible phase indicate that a change in stimulus can be detected using MVPA. Importantly, the results from the occluded phase indicate, that this effect is not due to the change in stimulus properties but due to the expectation violation as no change in speed was observable there. This pattern of results also suggests that the observed effects in V1 might be at least partially due to feedback, as they depended on the reappearance of the stimulus. Importantly, this region of V1 which encoded the trajectory of the stimulus after reappearance was not included in the analysis, thus any observed effects must be due to feedback mechanisms, for instance from higher visual areas (e.g. V5 Alink et al., 2010; Noesselt et al, 2002), frontal areas (Summerfield et al.,2006), or even hippocampal regions (Ekman et al., 2023). Moreover, our findings are in line with a recent study in mice which also investigated expectation violation in V1 (Tang et al, 2023). The study revealed that when a stimulus violated the mice’s expectation, neural response of V1 were enhanced, leading to a better encoding of the sensory information. The effect could be seen in both firing rate and neuronal synchrony in V1.

Future fMRI-studies with increased resolution could disentangle the effects of congruent and incongruent trials by segregating the areas of V1 which represent the horizontal and vertical trajectory in greater detail and would thus be able to separately test the similarity of responses for fast and slow velocities in congruent and incongruent trials. Moreover, the effect of attention in motion extrapolation needs to be tested by parametrically modulating attentional demands and compared whether the effect remains unchanged.

In conclusion, we tested whether visible motion processing or motion prediction would differentially modulate fMRI-responses in the primary visual cortex in higher or lower predictability contexts. According to the predictive coding model, it was expected that stimulation presented in low predictable context would enhance activity in V1 compared to high predictable contexts. However, alternatively, activity during high predictable context would also increase activity in this region if any amount of attention outweighs predictive mechanism in an all-or-nothing rather than a gradual way. Our results provided evidence supporting the latter hypothesis, by showing no difference in fMRI-response signal of high and low predictable context tasks when analysis the effects of accumulated evidence. When analysing short term adaptation effects, our trial-history analysis pointed to differences between consecutive trials of different categories compared to same category, suggesting that volatility at the trial-level may have increased activity in V1 and might thus be the more relevant time scale when analysing effects in V1. This difference was further supported by our MVPA results which showed difference in the decoding of the representational pattern of congruent and incongruent trials.

## Acknowledgement

We thank Denise Scheermann and Claus Tempelmann (Department of Neurology, Otto-von-Guericke Universität) for their support with scanning

## Ethics statement

This study was approved by the ethics committee of the Otto-von-Guericke-University.

## Funding statetement

CSA was funded by International Graduate School ABINEP (Analysis, Imaging, and Modelling of Neuronal and Inflammatory Processes) at Otto von Guericke University (OVGU) Magdeburg, Germany, kindly supported by the European Structural and Investment Funds (ESF, 2014–2020). TN was supported by DFG-SFB1436/B06. HH was supported by the “Autonomie im Alter” research consortium funded by the State of Saxony-Anhalt and the European Union, European Regional Development Fund (ERDF).

## Data availability statement

Data will be shared upon request with the need for approval from the requesting researcher’s local ethics committee.

## Footnotes

1 The first subjects had few volumes less due to the adjustment of scanning time and end of the experiment. One participant had 8 volumes less and four participants had 5 volumes less in the end of each run.

## REFERENCES

Abdollahi, R. O., Kolster, H., Glasser, M. F., Robinson, E. C., Coalson, T. S., Dierker, D., Jenkinson, M., Van Essen, D. C., & Orban, G. A. (2014). Correspondences between retinotopic areas and myelin maps in human visual cortex. NeuroImage, 99(100), 509–524. 10.1016/j.neuroimage.2014.06.042.

Agostino, C. S., Merkel, C., Ball, F., Vavra, P., Hinrichs, H., & Noesselt, T. (2023). Seeing and extrapolating motion trajectories share common informative activation patterns in primary visual cortex. Human Brain Mapping, 44(4), 1389–1406. 10.1002/hbm.26123

Aitchison, L., & Lengyel, M. (2017). With or without you: predictive coding and Bayesian inference in the brain. Current opinion in neurobiology, 46, 219–227. 10.1016/j.conb.2017.08.010

Alink, A., Schwiedrzik, C. M., Kohler, A., Singer, W., & Muckli, L. (2010). Stimulus predictability reduces responses in primary visual cortex. The Journal of neuroscience: the official journal of the Society for Neuroscience, 30(8), 2960–2966. 10.1523/JNEUROSCI.3730-10.2010

Baess, P., Widmann, A., Roye, A., Schröger, E., & Jacobsen, T. (2009). Attenuated human auditory middle latency response and evoked 40-Hz response to self-initiated sounds. The European journal of neuroscience, 29(7), 1514–1521. 10.1111/j.1460-9568.2009.06683.x

Battaglini, L., & Ghiani, A. (2021). Motion behind occluder: Amodal perception and visual motion extrapolation. Visual Cognition, 29(8), 475–499. 10.1080/13506285.2021.1943094

Behrens, T. E., Woolrich, M. W., Walton, M. E., & Rushworth, M. F. (2007). Learning the value of information in an uncertain world. Nature neuroscience, 10(9), 1214–1221. 10.1038/nn1954

Bordier, C., Hupé, J. M., & Dojat, M. (2015). Quantitative evaluation of fMRI retinotopic maps, from V1 to V4, for cognitive experiments. Frontiers in human neuroscience, 9, 277. 10.3389/fnhum.2015.00277.

Brett. M., Anton J.L., Valabregue, R., Poline, J.B. (2002). Region of interest analysis using an SPM toolbox [abstract] presented at the 8th International Conference on Functional Mapping of the Human Brain, June 2-6, 2002, Sendai, Japan. Available on CD-ROM in NeuroImage, Vol 16, No 2.

Carrasco, M., Talgar, C. P., & Cameron, E. L. (2001). Characterizing visual performance fields: effects of transient covert attention, spatial frequency, eccentricity, task and set size. Spatial vision, 15(1), 61–75. 10.1163/15685680152692015.

Caudek, C., Ceccarini, F., & Sica, C. (2015). Posterror speeding after threat-detection failure. Journal of experimental psychology. Human perception and performance, 41(2), 324–341. 10.1037/a0038753

Coull, J. T., Vidal, F., Goulon, C., Nazarian, B., & Craig, C. (2008). Using time-to-contact information to assess potential collision modulates both visual and temporal prediction networks. Frontiers in human neuroscience, 2, 10. 10.3389/neuro.09.010.2008

Dale, A. M., & Buckner, R. L. (1997). Selective averaging of rapidly presented individual trials using fMRI. Human Brain Mapping, 5(5), 329–340.

Danielmeier, C., & Ullsperger, M. (2011). Post-error adjustments. Frontiers in psychology, 2, 233. 10.3389/fpsyg.2011.00233

Di Russo, F., Martínez, A., Sereno, M. I., Pitzalis, S., & Hillyard, S. A. (2002). Cortical sources of the early components of the visual evoked potential. Human brain mapping, 15(2), 95–111. 10.1002/hbm.10010

Doherty, J. R., Rao, A., Mesulam, M. M., & Nobre, A. C. (2005). Synergistic effect of combined temporal and spatial expectations on visual attention. The Journal of neuroscience: the official journal of the Society for Neuroscience, 25(36), 8259–8266. 10.1523/JNEUROSCI.1821-05.2005

Edwards G, Vetter P, McGruer F, Petro LS, Muckli L. (2017). Predictive feedback to V1 dynamically updates with sensory input. Sci Rep, 28, 7(1):16538. doi: 10.1038/s41598-017-16093-y

Ekman, M., Kok, P., & de Lange, F. P. (2017). Time-compressed preplay of anticipated events in human primary visual cortex. Nature communications, 8, 15276. 10.1038/ncomms15276.

Ekman, M., Kusch, S., & de Lange, F. P. (2023). Successor-like representation guides the prediction of future events in human visual cortex and hippocampus. eLife, 12, e78904. 10.7554/eLife.78904

Etzel, J. A. (2017). MVPA significance testing when just above chance, and related properties of permutation tests. In 2017 International Workshop on Pattern Recognition in Neuroimaging (PRNI) (pp. 1–4). IEEE. DOI: 10.1109/PRNI.2017.7981498

Etzel, J. A. and Braver, T. S. (2013). “MVPA Permutation Schemes: Permutation Testing in the Land of Cross-Validation”. International Workshop on Pattern Recognition in Neuroimaging, 2013, pp. 140–143, doi: 10.1109/PRNI.2013.44.

Fischer, R., Plessow, F., & Ruge, H. (2013). Priming of visual cortex by temporal attention? The effects of temporal predictability on stimulus(-specific) processing in early visual cortical areas. NeuroImage, 66, 261–269. 10.1016/j.neuroimage.2012.10.091

Foxe, J. J., Murray, M. M., & Javitt, D. C. (2005). Filling-in in schizophrenia: a high-density electrical mapping and source-analysis investigation of illusory contour processing. Cerebral cortex (New York, N.Y. : 1991), 15(12), 1914–1927. 10.1093/cercor/bhi069

Friston K. (2003). Learning and inference in the brain. Neural networks: the official journal of the International Neural Network Society, 16(9), 1325–1352. 10.1016/j.neunet.2003.06.005

Friston, K. J., Fletcher, P., Josephs, O., Holmes, A., Rugg, M. D., & Turner, R. (1998). Event-related fMRI: characterizing differential responses. NeuroImage, 7(1), 30–40. 10.1006/nimg.1997.0306

Friston, K. J., Holmes, A. P., Poline, J. B., Grasby, P. J., Williams, S. C., Frackowiak, R. S., & Turner, R. (1995). Analysis of fMRI time-series revisited. NeuroImage, 2(1), 45–53. 10.1006/nimg.1995.1007

Georgieva, S., Peeters, R., Kolster, H., Todd, J. T., & Orban, G. A. (2009). The processing of three-dimensional shape from disparity in the human brain. The Journal of neuroscience : the official journal of the Society for Neuroscience, 29(3), 727–742. 10.1523/JNEUROSCI.4753-08.2009.

Heilbron, M., & Chait, M. (2018). Great Expectations: Is there Evidence for Predictive Coding in Auditory Cortex?. Neuroscience, 389, 54–73. 10.1016/j.neuroscience.2017.07.061

Helmholtz, H. (1866/1962). Concerning the perceptions in general. In J. Southall (Ed.) Treatise on physiological optics (3rd ed., Vol. III). New York: Dover, (Translation).

Hinrichs, H., Scholz, M., Tempelmann, C., Woldorff, M. G., Dale, A. M., & Heinze, H. J. (2000). Deconvolution of event-related fMRI responses in fast-rate experimental designs: tracking amplitude variations. Journal of cognitive neuroscience, 12 Suppl 2, 76–89. 10.1162/089892900564082

Johnson, P. A., Blom, T., van Gaal, S., Feuerriegel, D., Bode, S., & Hogendoorn, H. (2023). Position representations of moving objects align with real-time position in the early visual response. eLife, 12, e82424.

Kahneman, D., & Tversky, A. (1972). Subjective probability: A judgment of representativeness. Cognitive Psychology, 3(3), 430–454. 10.1016/0010-0285(72)90016-3

Karim, A. K., & Kojima, H. (2010). The what and why of perceptual asymmetries in the visual domain. Advances in cognitive psychology, 6, 103–115. 10.2478/v10053-008-0080-6.

King, J. A., Korb, F. M., von Cramon, D. Y., & Ullsperger, M. (2010). Post-error behavioral adjustments are facilitated by activation and suppression of task-relevant and task-irrelevant information processing. The Journal of neuroscience: the official journal of the Society for Neuroscience, 30(38), 12759–12769. 10.1523/JNEUROSCI.3274-10.2010

Kok, P., Rahnev, D., Jehee, J. F., Lau, H. C., & de Lange, F. P. (2012). Attention reverses the effect of prediction in silencing sensory signals. Cerebral cortex (New York, N.Y. : 1991), 22(9), 2197–2206. 10.1093/cercor/bhr310

Kolster, H., Peeters, R., & Orban, G. A. (2010). The retinotopic organization of the human middle temporal area MT/V5 and its cortical neighbors. The Journal of neuroscience: the official journal of the Society for Neuroscience, 30(29), 9801–9820. 10.1523/JNEUROSCI.2069-10.2010

Krala, M., van Kemenade, B., Straube, B., Kircher, T., & Bremmer, F. (2019). Predictive coding in a multisensory path integration task: An fMRI study. Journal of vision, 19(11), 13. 10.1167/19.11.13.

Lakha, L., & Humphreys, G. (2005). Lower visual field advantage for motion segmentation during high competition for selection. Spatial vision, 18(4), 447–460. 10.1163/1568568054389570.

Levine, M. W., & McAnany, J. J. (2005). The relative capabilities of the upper and lower visual hemifields. Vision research, 45(21), 2820–2830. 10.1016/j.visres.2005.04.001.

Muckli L, Kohler A, Kriegeskorte N, Singer W. (2005). Primary visual cortex activity along the apparent-motion trace reflects illusory perception. PLoS Biol, 3(8):e265. doi: 10.1371/journal.pbio.0030265.

Mumford D. (1992). On the computational architecture of the neocortex. II. The role of cortico-cortical loops. Biological cybernetics, 66(3), 241–251. 10.1007/BF00198477

Mumford, J. A., Turner, B. O., Ashby, F. G., & Poldrack, R. A. (2012). Deconvolving BOLD activation in event-related designs for multivoxel pattern classification analyses. NeuroImage, 59(3), 2636–2643. 10.1016/j.neuroimage.2011.08.076

Nasr, S., & Tootell, R. (2018). Visual field biases for near and far stimuli in disparity selective columns in human visual cortex. NeuroImage, 168, 358–365. 10.1016/j.neuroimage.2016.09.012.

Nasr, S., & Tootell, R. (2020). Asymmetries in Global Perception Are Represented in Near-versus Far-Preferring Clusters in Human Visual Cortex. The Journal of neuroscience : the official journal of the Society for Neuroscience, 40(2), 355–368. 10.1523/JNEUROSCI.2124-19.2019.

Noesselt, T., Hillyard, S. A., Woldorff, M. G., Schoenfeld, A., Hagner, T., Jäncke, L., Tempelmann, C., Hinrichs, H., & Heinze, H. J. (2002). Delayed striate cortical activation during spatial attention. Neuron, 35(3), 575–587. 10.1016/s0896-6273(02)00781-x

Oosterhof, N. N., Connolly, A. C., and Haxby, J. V. (2016). CoSMoMVPA: multi-modal multivariate pattern analysis of neuroimaging data in Matlab / GNU Octave. Frontiers in Neuroinformatics, doi:10.3389/fninf.2016.00027.

Patton, Lydia, “Hermann von Helmholtz”, The Stanford Encyclopedia of Philosophy (Winter 2018 Edition), Edward N. Zalta (ed.), URL = <https://plato.stanford.edu/archives/win2018/entries/hermann-helmholtz/>

Previc, F. (1990). Functional specialization in the lower and upper visual fields in humans: Its ecological origins and neurophysiological implications. Behavioral and Brain Sciences, 13(3), 519–542. doi:10.1017/S0140525X00080018

Rao, R. P., & Ballard, D. H. (1999). Predictive coding in the visual cortex: a functional interpretation of some extra-classical receptive-field effects. Nature neuroscience, 2(1), 79–87. 10.1038/4580

Rijsdijk, J. P., Kroon, J. N., & van der Wildt, G. J. (1980). Contrast sensitivity as a function of position on the retina. Vision research, 20(3), 235–241. 10.1016/0042-6989(80)90108-x

Rissman, J., Gazzaley, A., & D’Esposito, M. (2004). Measuring functional connectivity during distinct stages of a cognitive task. NeuroImage, 23(2), 752–763.

Schellekens, W., van Wezel, R. J., Petridou, N., Ramsey, N. F., & Raemaekers, M. (2016). Predictive coding for motion stimuli in human early visual cortex. Brain structure & function, 221(2), 879–890. 10.1007/s00429-014-0942-2

Shipp, S., Adams, R. A., & Friston, K. J. (2013). Reflections on agranular architecture: predictive coding in the motor cortex. Trends in neurosciences, 36(12), 706–716. 10.1016/j.tins.2013.09.004

Skrandies, W. (1987). The Upper and Lower Visual Field of Man: Electrophysiological and Functional Differences. In: Autrum, H., Ottoson, D., Perl, E.R., Schmidt, R.F., Shimazu, H., Willis, W.D. (eds) Progress in Sensory Physiology. Progress in Sensory Physiology, vol 8. Springer, Berlin, Heidelberg. 10.1007/978-3-642-71060-5_1

Spratling M. W. (2017). A review of predictive coding algorithms. Brain and cognition, 112, 92–97. 10.1016/j.bandc.2015.11.003

Sterzer P, Haynes JD, Rees G. (2006). Primary visual cortex activation on the path of apparent motion is mediated by feedback from hMT+/V5. Neuroimage, 32(3):1308–16. doi: 10.1016/j.neuroimage.2006.05.029.

Summerfield, C., Egner, T., Greene, M., Koechlin, E., Mangels, J., & Hirsch, J. (2006). Predictive codes for forthcoming perception in the frontal cortex. Science (New York N.Y.), 314(5803), 1311–1314. 10.1126/science.1132028

Talgar, C. P., & Carrasco, M. (2002). Vertical meridian asymmetry in spatial resolution: visual and attentional factors. Psychonomic bulletin & review, 9(4), 714–722. 10.3758/bf03196326

Tang, M. F., Kheradpezhouh, E., Lee, C. C., Dickinson, J. E., Mattingley, J. B., & Arabzadeh, E. (2023). Expectation violations enhance neuronal encoding of sensory information in mouse primary visual cortex. Nature Communications, 14(1), 1196.

Tversky, A., & Kahneman, D. (1971). Belief in the law of small numbers. Psychological Bulletin, 76(2), 105–110. 10.1037/h0031322

Tversky, A., & Kahneman, D. (1973). Availability: A heuristic for judging frequency and probability. Cognitive psychology, 5(2), 207–232. 10.1016/0010-0285(73)90033-9

Vetter, P., Grosbras, M. H., & Muckli, L. (2015). TMS over V5 disrupts motion prediction. Cerebral cortex (New York, N.Y. : 1991), 25(4), 1052–1059. 10.1093/cercor/bht297

Warnking, J., Dojat, M., Guérin-Dugué, A., Delon-Martin, C., Olympieff, S., Richard, N., Chéhikian, A., & Segebarth, C. (2002). fMRI retinotopic mapping--step by step. NeuroImage, 17(4), 1665–1683. 10.1006/nimg.2002.1304.

Zylberberg, A., Fetsch, C. R., & Shadlen, M. N. (2016). The influence of evidence volatility on choice, reaction time and confidence in a perceptual decision. eLife, 5, e17688. 10.7554/eLife.17688

