## Supplementary Material for "Contextual modulation of primary visual cortex by temporal predictability during motion extrapolation"

### 1.Results

#### 1.1. Behaviour

##### 1.1.1. Temporal Estimation

*Visible Phase*

*Low predictable context*: We further observed an interaction between velocity and congruency (F(1,14)=52.895, p<.001, $ŋ_p^2$=.791; congruent fast vs. incongruent fast: MD=-0.089, ±.011, t=-8.292, $p_{bonf}$ <.001, and congruent slow vs. incongruent slow: MD=.040, ±.011, t=3.709, $p_{bonf}$ =.006). Difference between incongruent fast and slow was significantly high (MD=.080, ±.010, t=8.072, $p_{bonf}$<.001). Main effects were seen for congruency (F(1,14)=16.446, p=.001, $ŋ_p^2$=.540, congruent-incongruent: MD=-0.025, ±.006, t=-4.055, $p_{bonf}$=.001) and velocity(F(1,14)=12.269, p=.004, $ŋ_p^2$=.467, fast-slow: MD=.015, ±.004, t=3.503, $p_{bonf}$=.004).

*High x Low Predictability*: Significant differences were observed in HP context between upwards fast and slow motion (MD=-0.091, ±.013, t=-7.087, $p_{bonf}$<.001) and downward fast and slow motion (MD=-0.114, ±.013, t=-8.892, $p_{bonf}$<.001). However, differences between downward and upward trajectories were also observed during slow motion (MD=-0.043, ±.012, t=-3.540, $p_{bonf}$=.027). In LP context, differences were seen between upward fast and slow motion (MD= -0.065, ±.013, t=-5.065, p<.001). Reaction time was also significantly different in HP compared to LP context during downward slow motion (MD= .073, ±.011, t=6.457, $p_{bonf}$<.001). We further observed an interaction between predictability and velocity (F(1,14)=28.829, $p_{bonf}$<.001, $ŋ_p^2$ =.673), with differences between fast and slow motion in HP context (fast-slow: MD=-0.103, ±.008, t=-13.345, $p_{bonf}$<.001); in LP context (fast-slow: MD=-0.049, ±.008, t=-6.409, $p_{bonf}$<.001), and between HP and LP context, indicating higher reaction time during slow motion in HP (fast-slow: MD=.039, ±.009, t=4.501, $p_{bonf}$<.001). An interaction between predictability and direction (F(1,14)=23.795, p<.001, $ŋ_p^2$ =.630) pointed to a significant difference in HP context between upward and downward direction (MD=-0.032, ±.007, t=-4.787, $p_{bonf}$<.001); and between HP and LP during downward direction (MD=.033, ±.008, t=3.956, $p_{bonf}$=.004). Additionally, we observed a main effect of velocity (F(1,14)=167.373, p<.001, $ŋ_p^2$ =.923; fast vs. slow: MD=-0.076, ±.006, t=-12.937, $p_{bonf}$<.001) and direction (F(1,14)=4.996, p=.042, $ŋ_p^2$ =.263; upward vs. downward: MD=-0.012, ±.005, t=-2.235, $p_{bonf}$=.042).

*Occluded Phase*

*High x Low Predictability:* We also observed significant main effects for direction ($F(1,14)=13.216$, $p=.003$, $\eta_p^2=.486$; upward vs. downward: MD=-0.012, ±.003, t=-3.635, $p_{bonf}=.003$), velocity ($F(1,14)=45.490$, $p<.001$, $\eta_p^2=.765$; fast vs. slow: MD=-0.057, ±.008, t=-6.745, $p_{bonf}<.001$) and a marginally significant effect for predictability ($F(1,14)=4.136$, $p=.061$, $\eta_p^2=.228$; HP vs. LP context: MD=-0.024, ±.012, t=2.034, $p_{bonf}=.061$). direction and predictability ($F(1,14)=10.252$, $p=.006$, $\eta_p^2=.423$; upward vs. downward in HP context: MD=-0.023, ±.005, t=-4.822, $p_{bonf}<.001$). Interaction between direction and velocity did not reach significance, but post-hocs show significant differences between conditions, indicating that participants were estimating time-to-contact accordingly.

##### 1.1.2. Temporal Estimation Error

*Visible Phase*

*High predictable context*: Results showed main effect of velocity ($F(1,14)=137.095$, $p<.001$, $\eta_p^2=.907$; fast vs. slow: (MD=.098, ±.008, t=11.709, $p_{bonf}<.001$) and direction ($F(1,14)=16.309$, $p=.001$, $\eta_p^2$ .538; upward vs. downward: MD=-0.032, ±.008, t=-4.038, $p_{bonf}=.001$), revealing an overestimation for fast motion and upward direction. Analysis did not reveal significant interaction. Figure-supp. 1A depicts averaged reaction time error for all conditions during visible phase.

*Low predictable context*: We observed again the triple interaction between direction, velocity and congruency (Figure-supp. 1A), suggesting that errors were smaller during fast incongruent trials than during congruent trials in upward direction, but not downward direction ($F(1,14)=12.269$, $p=.004$, $\eta_p^2=.467$; congruent fast vs. incongruent fast in upward: MD=.159, ±.012, t=13.660, $p<.001$; congruent fast vs. incongruent fast in downward: MD=.125, ±.012, t=10.686, $p<.001$); congruent slow vs. incongruent slow in upward (MD=-0.142, ±.012, t=-9.904, $p<.001$; congruent slow vs. incongruent slow in downward: MD=-0.115, ±.012, t=10.686, $p<.001$). Results still revealed an interaction between velocity and congruency ($F(1,14)=231.417$, $p<.001$, $\eta_p^2=.943$; congruent fast vs. incongruent fast: MD=.142, ±.010, t=14.611, $p_{bonf}=.001$; congruent slow vs. incongruent slow: MD=-0.129, ±-0.129, ±.010, t=-13.259, $p_{bonf}<.001$). Main effects were observed for direction ($F(1,14)=16.445$, $p=.001$, $\eta_p^2=.540$, up vs. down: MD=-0.019, ±.008, t=-2.425, $p_{bonf}<.001$), and velocity ($F(1,14)=5.882$, $p=.029$, $\eta_p^2=.296$; fast vs. slow: MD=-0.019, ±.008, $p_{bonf}=.029$).

*High x Low Predictability*: Analysis of congruent trials of HP and LP contexts from pointed to an interaction between predictability and velocity ($F(1,14)=4.930$, $P=.043$, $\eta_P^2=.260$), similar to results from reaction time. Differences were observed between fast and slow motion in HP (MD=.097, ±.011, t=8.881, $p_{bonf}<.001$) and LP (MD=.117, ±.011, t=-10.663, $p_{bonf}<.001$). Main effects were observed for direction ($F(1,14)=10.229$, $p=.006$, $\eta_p^2=.424$; upward vs. downward: MD=-0.027, ±.008, t=-3.209, $p_{bonf}=.006$) and velocity ($F(1,14)=113.799$, $p<.001$, $\eta_p^2=.860$; fast vs. slow: MD=.107, ±.010, t=10.668, $p_{bonf}<.001$).

<u>*Occluded Phase*</u>

*High Predictable Context*: Main effect was observed for velocity ($F(1,14)=44.235$, $p<.001$, $\eta_p^2=.747$; fast vs. slow: MD=-0.086, ±.013. t=-6.651, $p_{bonf}<.001$) and direction ($F(1,14)=16.420$, $p=.001$, $\eta_p^2=.523$; up vs. down: MD=-0.024, ±.006. t=-4.052, $p_{bonf}=.001$), showing an overestimation, this time, during fast motion as well as during downward direction. No significant interactions were observed. Figure-supp. 1B depicts averaged reaction time error for all conditions during occluded phase.

*Low Predictable Context*: We observed an interaction between velocity and congruency, indicating an overestimation during congruent fast and incongruent slow (speeded-up stimulus)($F(1,14)=483.345$, $p<.001$, $\eta_p^2=.972$; congruent fast vs. congruent slow: MD=.0172, ±.014, t=11.879, $p_{bonf}<.001$; congruent fast vs. incongruent fast: MD=.193, ±.010, t=19.806, $p_{bonf}<.001$; congruent slow vs. incongruent slow: MD=-0.190, ±.010, t=-19.525, $p_{bonf}<.001$; incongruent fast vs. incongruent slow: MD=-0.212, ±.014, t=-14.658, $p_{bonf}<.001$). A marginally significant triple interaction (Figure-supp. 1B) was observed (direction, velocity and congruency: $F(1,14)=4.248$ , $p=.058$, $\eta_p^2=.233$). Post-hoc comparisons pointed to differences between congruent fast vs. congruent slow in upward (MD=.168, ±.019, t=8.904, $p_{bonf}<.001$); congruent fast vs. congruent slow in downward (MD=.175, ±.019, t=9.260, $p_{bonf}<.001$); congruent fast vs. incongruent fast in upward (MD=.198, ±.011, t=17.939, $p_{bonf}<.001$); congruent slow vs. incongruent slow in upward (MD=-0.200, ±.011, t=-18.065, $p_{bonf}<.001$); congruent fast vs. incongruent fast in downward (MD=.188, ±.011, t=17.009, $p_{bonf}<.001$); incongruent fast vs. incongruent slow in upward (MD=-0.230, ±.019, t=-12.148, $p_{bonf}<.001$); and incongruent fast vs. incongruent slow in downward (MD=-0.194, ±.019, t=-10.299, $p_{bonf}<.001$).

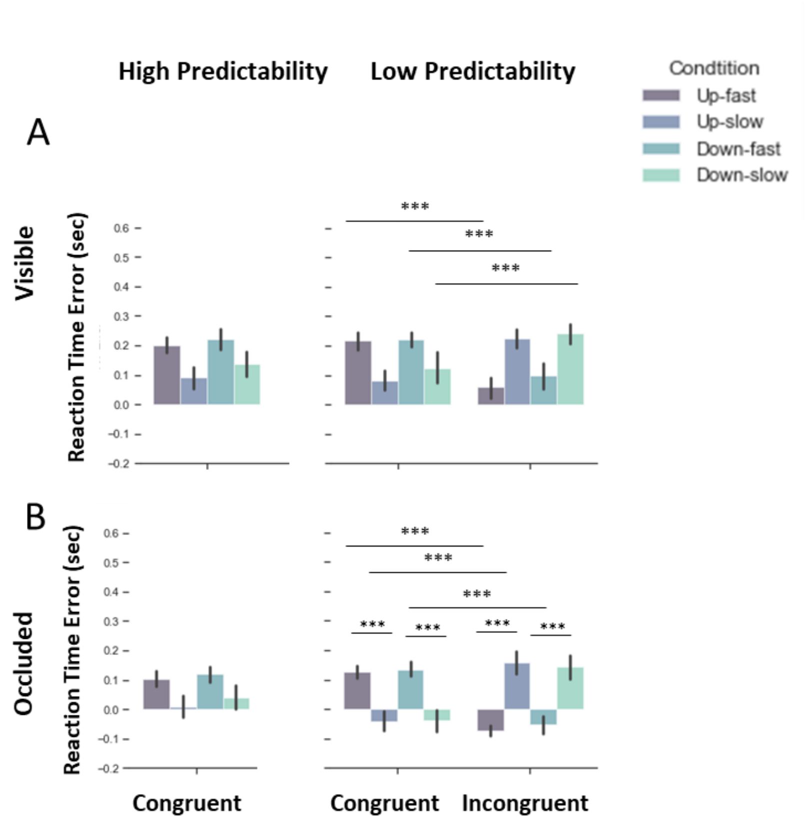


*Figure-supp. 1- Averaged reaction time error across conditions. Dark purple bars depict results of fast motion in upward trajectory, blue bars depict results of slow motion in upward trajectory, dark green bar, fast motion in downward direction and light green, slow motion in downward trajectory. Although the conditions were clearly (and significantly) different in HP context during visible (A-left) phase and occluded (B-left), the analysis did not show interaction between direction and velocity. (A) Differences were observed in LP context during visible phase and occluded phases (B) when congruent and incongruent trials were compared. Note that the bars in fast congruent conditions are similar to the slow in incongruent conditions, showing that participants were trying to estimate according to the congruent velocity of the stimulus. During the slow conditions (and incongruent fast), we could observe an underestimation, while during fast (and incongruent slow) condition we see that participants overestimated their responses. In appendix B, it is possible to observe a plot of reaction time error of incongruent condition during occlusion in the LP context calculated based on the expected velocity subtracted from participants' response. Interestingly, we observed the same pattern of response seen in congruent condition during occlusion in the LP context, suggesting that participants were not estimating according to the feedback, rather they were responding according to the learned velocity-direction association.*

*High x Low Predictability*: Reaction time error analysis yielded similar results compared to reaction time. We observed an interaction for direction and predictability ($F(1,14)=5.069$, $p=.041$, $\eta_p^2=.266$; upward vs. downward in HP context: $MD=-0.023$, $\pm.006$, $t=-3.914$, $p_{bonf}=.003$), velocity and predictability ($F(1,14)=38.000$, $p<.001$, $\eta_p^2=.731$; fast vs. slow in HP context: MD=.088, ±.013, t=6.873, $p_{bonf}$<.001; slow in HP vs. slow in LP context: MD=.063, ±.013, t=4.890, $p_{bonf}$ <.001; fast vs. slow in LP context: MD=.172, ±.013, t=13.420, $p_{bonf}$ <.001). We also observed significant main effects for direction (F(1,14)=10.491, p=.006, $\eta_p^2$=.428; upward vs. downward: MD=-0.014, ±.004, t=-3.239, $p_{bonf}$=.006), velocity (F(1,14)=143.372, p<.001, $\eta_p^2$=.911; fast vs. slow: MD=.130, ±.011, t=11.974, $p_{bonf}$<.001) and a marginally significant effect for predictability (F(1,14)=3.783, p=.072, $\eta_p^2$ =.213).

##### 1.1.3. Spatial estimation

*Visible Phase*

*High predictable context*: Results indicated a main effect of velocity, suggesting a higher performance during slow motion (F(1,14)=44.061, p <.001, $\eta_p^2$=0.759; fast-slow: mean difference (MD)=-0.058, (SE)±.009, t=-6.638, $p_{bonf}$ <.001) and marginally significant main effect for direction, pointing to a tendency of higher performance during upward direction (F(1,14)=3.709, p=.075, $\eta_p^2$=.209; upward vs. downward: MD=.032, ±.016, t=2.926, $p_{bonf}$=.075). No interaction was observed between factors. Figure-supp. 2A shows averaged accuracy for all direction-velocity paired conditions during visible period.

*Low predictable context*: We observed a triple interaction (Figure-supp. 2A) between direction, velocity and congruency (F(1,14)=19.174, p<.001, $\eta_p^2$ =.578; congruent fast vs. incongruent fast in downward: MD=.104, ±.022, t=4.736, $p_{bonf}$<.001; congruent slow vs. incongruent slow in upward: MD=.102, ±.022, t=4.643, $p_{bonf}$<.001). Among incongruent conditions (incongruent fast=expected fast, but stimulus slowed down; incongruent slow=expected slow, but stimulus speeded up), we observed differences between both velocities in both upwards and downwards direction (upward fast vs. downward fast: MD=.108, ±.029, t=3.785, $p_{bonf}$=.012, and upward slow vs. downward slow: MD=-0.107, ±.029, t=-3.759, $p_{bonf}$=.013). Within downward direction, performance during incongruent slow motion was significantly higher (downward fast vs. downward slow: MD=-0.128, ±.026, t=-4.871, $p_{bonf}$<.001). We still found an interaction between direction and velocity (F(1,14)=5.014, p=.042, $\eta_p^2$ =.264; fast vs. slow in downward: MD=-0.068, ±.021, t=-3.219, $p_{bonf}$=0.024). Additionally, results indicated main effect of congruency (F(1,14)=10.721, p=.006, $\eta_p^2$ =.434, congruent vs. incongruent: MD=.037, ±.011, t=3.274, $p_{bonf}$=.006) and a main effect of velocity (F(1,14)=6.914, p=.020, $\eta_p^2$ =.331, fast-slow: MD=-.026, ±.010, t=-2.630, $p_{bonf}$=.020).

*High x Low Predictability*: In the comparison of congruent results from both high and low predictable phases during the visible task, we observed a triple interaction between direction, velocity and predictability (F(1,14)=6.481, p=.023, $\eta_p^2$ =.316). Significant differences were observed in LP context between upwards fast and slow motion (MD=-0.056, ±.016, t=-3.498, p=.028); in HP context between downward fast and slow motion (MD= -0.077, ±.016, t=-4.777, p<.001) and between HP and LP contexts during downward fast motion (MD= -0.059, ±.015, t=-3.829, p=.010). Results also pointed to interactions between predictability and velocity (F(1,14)=6.800, p=.021, $\eta_p^2$ =.327), with differences between fast and slow motion in HP context (fast vs. slow: MD=-0.058, ±.010, t=-5.945, p<.001) and LP context (fast vs. slow: MD=-0.032, ±.010, t=-3.278, p=.020), indicating higher performance during slow motion. An interaction between predictability and direction (F(1,14)=7.167, p=.018, $\eta_p^2$ =.339) pointed to a marginally significant difference in HP context between upward and downward conditions (MD=.034, ±.013, t=2.574, p=.094). Finally, we observed a main effect of velocity (F(1,14)=28.794, p<.001, $\eta_p^2$ =.673; fast vs. slow: MD=-0.045, ±.008, t=-5.366, p<.001).

*Occluded Phase*

Please note that the pattern of correct response is almost identical for occluded and visible stimulation; thus, it might reflect participants' overall attentiveness.

*High Predictable Context*: As in the HP visible phase, here we observed only a main effect of velocity, again revealing higher performance during slow motion (F(1,14)= 15.996, p<.001, $\eta_p^2$ =.533; fast vs. slow: MD=-0.050, ±.013. t=-4, $p_{bonf}$<.001). No interaction was observed. Figure-supp. 2B shows averaged accuracy for all direction-velocity paired conditions during occlusion period.

*Low Predictable Context*: During this phase, analyses pointed to a triple interaction between direction, velocity and congruency (Figure-supp. 2B), indicating that higher performance during congruent trials compared to incongruent trials (F(1,14)= 23.749 p<.001, $\eta_p^2$ =.629; congruent fast vs. incongruent fast in downward: MD=.124, ±.024, t=5.140, $p_{bonf}$ <.001; congruent slow vs. incongruent slow in upward: MD=.100, ±.024, t=4.162, $p_{bonf}$ =.004). For incongruent trials different pattern of results were observed (incongruent fast in upwards vs. incongruent fast in downward: MD=.130, ±.028, t=4.670, $p_{bonf}$ =.001; incongruent fast vs. incongruent slow in upward: MD=.115, ±.029, t=3.910, $p_{bonf}$ =.009; incongruent fast vs. incongruent slow in downward: MD=-0.127, ±.029, t=-4.340, $p_{bonf}$=.002; incongruent slow in upwards vs. incongruent slow in downward: MD=-0.112, ±.028, t=-4.028, $p_{bonf}$=.007). An interaction between direction and velocity (F(1,14)= 4.731 p=.047, $\eta_p^2$ =.253; post-hoc failed to show significant results). A main effect of congruency was also revealed by the analysis (F(1,14)= 7.856 p =.014, $\eta_p^2$ =.359; congruent-incongruent: MD=.034, ±.012, t=2.803, $p_{bonf}$ =.014).

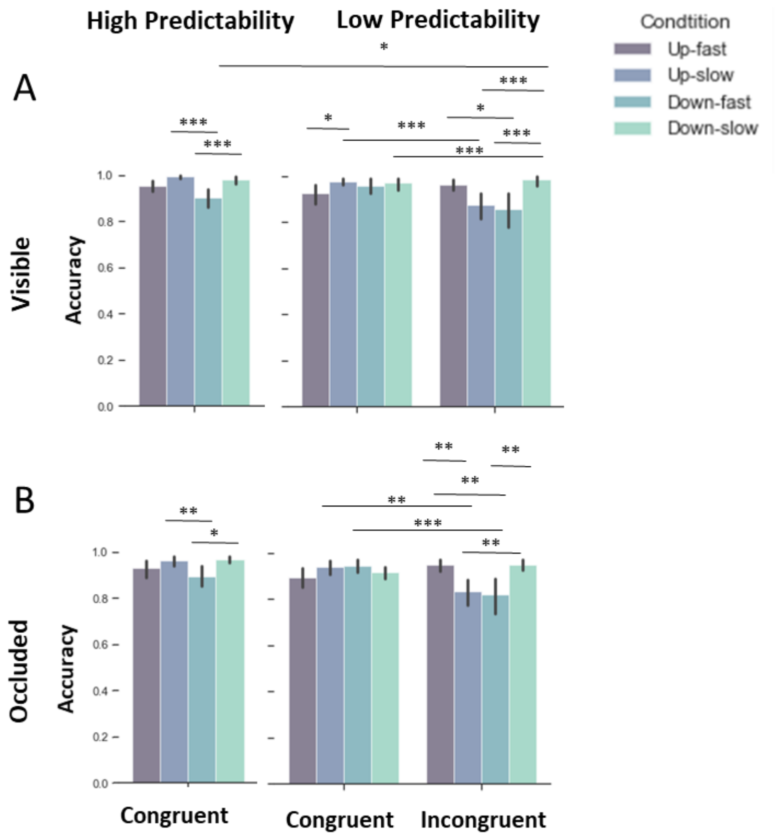


*Figure-supp. 2 - Averaged accuracy across conditions. Dark purple bars depict results of fast motion in upward trajectory, blue bars depict results of slow motion in upward trajectory, dark green bar, fast motion in downward direction and light green, slow motion in downward trajectory. Note that the accuracy results were highly similar for occluded and visible motion trajectories. (A) Results of visible period during HP (left) and LP contexts (right) show differences between conditions. Differences were observed between congruent fast and slow in downward condition in HP context, and also difference between the concurrent conditions up-slow/down-fast (condition presented within the same task). Analysis also yielded differences between congruent and incongruent conditions in LP contexts, as well as a significant difference between HP and LP, indicating that slow in downward trajectory in LP presented higher accuracy than slow during downward trajectory in HP. (B) Results of occluded period during HP (left) and LP contexts (right) showed differences between slow and fast in downward trajectory during HP, and also difference between the concurrent condition up-slow/down-fast (condition presented within the same task), as in visible phase. During occluded phase in LP contexts, differences between congruent and incongruent conditions were observed, as during the visible phase.*

*High x Low Predictability*: Here we compare the congruent results from both high and low predictable phases during the occluded task. Results indicated a triple interaction between direction, velocity and predictability, suggesting higher performance during slow motion in downward direction in HP context ($F(1,14)=16.085$, $p=.001$, $\eta_p^2=.535$; downward slow in LP context vs. downward slow in HP context: $MD=-0.052$, $\pm.015$, $t=-3.429$, $p_{bonf}=.035$; downward fast vs. downward slow in HP context: $MD=-0.070$, $\pm.019$, $t=-3.598$, $p_{bonf}=.021$). We also observed an interaction between velocity and predictability ($F(1,14)=18.557$, $p<.001$, $\eta_p^2=.570$; slow in LP vs. slow in HP context: $MD=-0.038$, $\pm.010$, $t=-3.849$, $p_{bonf}=.005$; fast vs. slow in HP context: $MD=-0.050$, $\pm.012$, $t=-4.213$, $p_{bonf}=.002$). Besides a main effect for velocity ($F(1,14)=6.684$, $p=.002$, $\eta_p^2=.323$; fast vs. slow: $MD=-0.028$, $\pm.011$, $t=-2.585$, $p_{bonf}=.022$) was observed, as well as a marginally significant main effect for predictability ($F(1,14)=3.337$, $p=.083$, $\eta_p^2=.192$).

##### 1.2. Univariate fMRI-results

*Visible Phase*

*Low predictable context*: Main effects for direction ($F(1,14)=5.841$, $p=.030$, $\eta_p^2=.294$; upward-downward: $MD=-0.663$, $\pm.274$, $t=-2.417$, $p_{bonf}=.030$), velocity ($F(1,14)=38.808$, $p<.001$, $\eta_p^2=.735$; fast-slow: $MD=1.727$, $\pm.277$, $t=6.230$, $p_{bonf}<.001$) were also observed.

##### 1.3. MVPA results

Table-supp. 1 shows results of decoding performed in hMT/V5+, following the same analyses carried out with V1 in the main text.

#### Temporal Information: Velocity

| | Predictability | Accuracy | SE | Permutation p-value | N spheres (average) | Paired T-Test |
|---|---|---|---|---|---|---|
| Visible-Occluded | High | 0.569 | 0.005 | <.001 | 81 (±18.38) | t=3.390. |
| | Low | 0.558 | 0.004 | <.001 | 73 (± 20.93) | p = .004 |
| Visible | High | 0.580 | 0.005 | <.001 | 73 (±20.16) | t=0.484. |
| | Low | 0.577 | 0.005 | <.001 | 58 (±20.87) | p = .636 |
| Occluded | High | 0.581 | 0.005 | <.001 | 76 (±18.42) | t=1.359. |
| | Low | 0.575 | 0.004 | <.001 | 67 (±18.63) | p = .194 |

#### Spatial Information: Direction

| | Predictability | Accuracy | SE | Permutation p-value | N spheres (average) | Paired T-Test |
|---|---|---|---|---|---|---|
| Visible-Occluded | High | 0.548 | 0.004 | <.001 | 66 (±21.61) | t=-5.628. |
| | Low | 0.571 | 0.005 | <.001 | 72 (±23.80) | p <.001 |
| Visible | High | 0.553 | 0.003 | <.001 | 49 (±24.03) | t=-5.378. |
| | Low | 0.581 | 0.005 | <.001 | 59 (±20.77) | p <.001 |
| Occluded | High | 0.558 | 0.004 | <.001 | 51 (±23.52) | t=-3.740. |
| | Low | 0.578 | 0.005 | <.001 | 58 (±21.17) | p = .002 |

*Table-supp. 1 – Decoding accuracy of temporal and spatial information: Accuracy values of all analyses of all conditions are displayed on the table above, together with the standard error of the mean (SE). Permutation p-values demonstrate that the permuted accuracy distributions were highly significant different from the true label accuracy distribution. The average number of spheres included in the sample of 5% most informative spheres can be seen in the range of 49 to 81.*
